# Semi-automated genome annotation using epigenomic data and Segway

**DOI:** 10.1101/080382

**Authors:** Eric G. Roberts, Mickaël Mendez, Coby Viner, Mehran Karimzadeh, Rachel Chan, Rachel Ancar, Davide Chicco, Jay R. Hesselberth, Anshul Kundaje, Michael M. Hoffman

**Affiliations:** Princess Margaret Cancer Centre, University of Toronto, Toronto, Ontario, Canada; Department of Computer Science, University of Toronto, Toronto, Ontario, Canada; Department of Medical Biophysics, University of Toronto, Toronto, Ontario, Canada; Engineering Physics Program, University of British Columbia, Vancouver, British Columbia, Canada; Department of Biochemistry and Molecular Genetics, University of Colorado Denver, Aurora, Colorado, USA; Department of Computer Science, Stanford University, Stanford, California, USA; Department of Genetics, Stanford University, Stanford, California, USA

## Abstract

Biochemical techniques measure many individual properties of chromatin along the genome. These properties include DNA accessibility (measured by DNase-seq) and the presence of individual transcription factors and histone modifications (measured by ChIP-seq). Segway is software that transforms multiple datasets on chromatin properties into a single annotation of the genome that a biologist can more easily interpret. This protocol describes how to use Segway to annotate the genome, starting with reads from a ChIP-seq experiment. It includes pre-processing of data, training the Segway model, annotating the genome, assigning biological meanings to labels, and visualizing the annotation in a genome browser.

## INTRODUCTION

Segway^1,2^ is software that discovers patterns in genomic signal datasets, and then transforms multiple datasets into a simple annotation, labeling the best pattern at every position in the genome. Each input data set comes from a biochemical technique that measures some property along the genome. Often the property relates to local chromatin biology, such as DNA accessibility (measured by DNase-seq^3,4^ or ATAC-seq^5^) and the presence of individual transcription factors and histone modifications (measured by ChIP-seq^6^). The input data could, however, include any property quantified along the genome in a locus-specific manner. Given some genome-aligned datasets, Segway constructs a statistical model of recurring patterns across these datasets. In the model, every base has a hidden *label* that determines which pattern is generated at that position. Then, Segway uses that model to annotate the whole genome automatically with the best label for every position. Finally, supporting tools visualize and summarize the model and annotation to reveal how these patterns associate with known and novel biological phenomena.

### Motivation

Researchers often have multiple functional genomic datasets that they wish to understand. While analysts have a rich choice of peak-calling methods^7,8,9^, post hoc comparisons of peak calls are unwieldy, at best. At worst, they have decreased power to detect phenomena associated with low signal in a single dataset that are revealed as significant when we jointly consider multiple datasets. To discover more potentially significant regions, one needs a method of integrative analysis across multiple datasets.

To perform integrative analysis on unprecedented quantities of genomic signal data, researchers in the ENCODE Pilot Project^10^ developed the first semi-automated genome annotation method, HMMSeg^10,11,12^. Semi-automated genome annotation jointly analyzes multiple datasets in an unsupervised fashion, allowing the discovery of both known and novel patterns. It usually works by creating a *segmentation*, which is an annotation that has one label at every position. Since the ENCODE Pilot, multiple semi-automated genome annotation methods have been developed^13,14,15,16,17,18^, including HMMSeg’s successor, Segway. Segway is one of the most powerful methods for semi-automated genome annotation methods, capable of analyzing multiple datasets at 1–base pair resolution, handling heterogeneous patterns of missing data, and modeling signal level directly rather than binarizing.

### Experimental design

In this protocol, we describe how to create a Segway annotation from ChIP-seq data (Figure 1). The process has the following major steps:

1. Create bedGraph^19^ signal from aligned reads from ChIP-seq data or download signal from an existing project.
2. Create Genomedata^20^ archives containing the signal data.
3. Train the Segway model.
4. Use Segway to produce an annotation in browser-extensible data (BED) format^21^.
5. View and analyze the resulting annotation.

**Figure 1:**
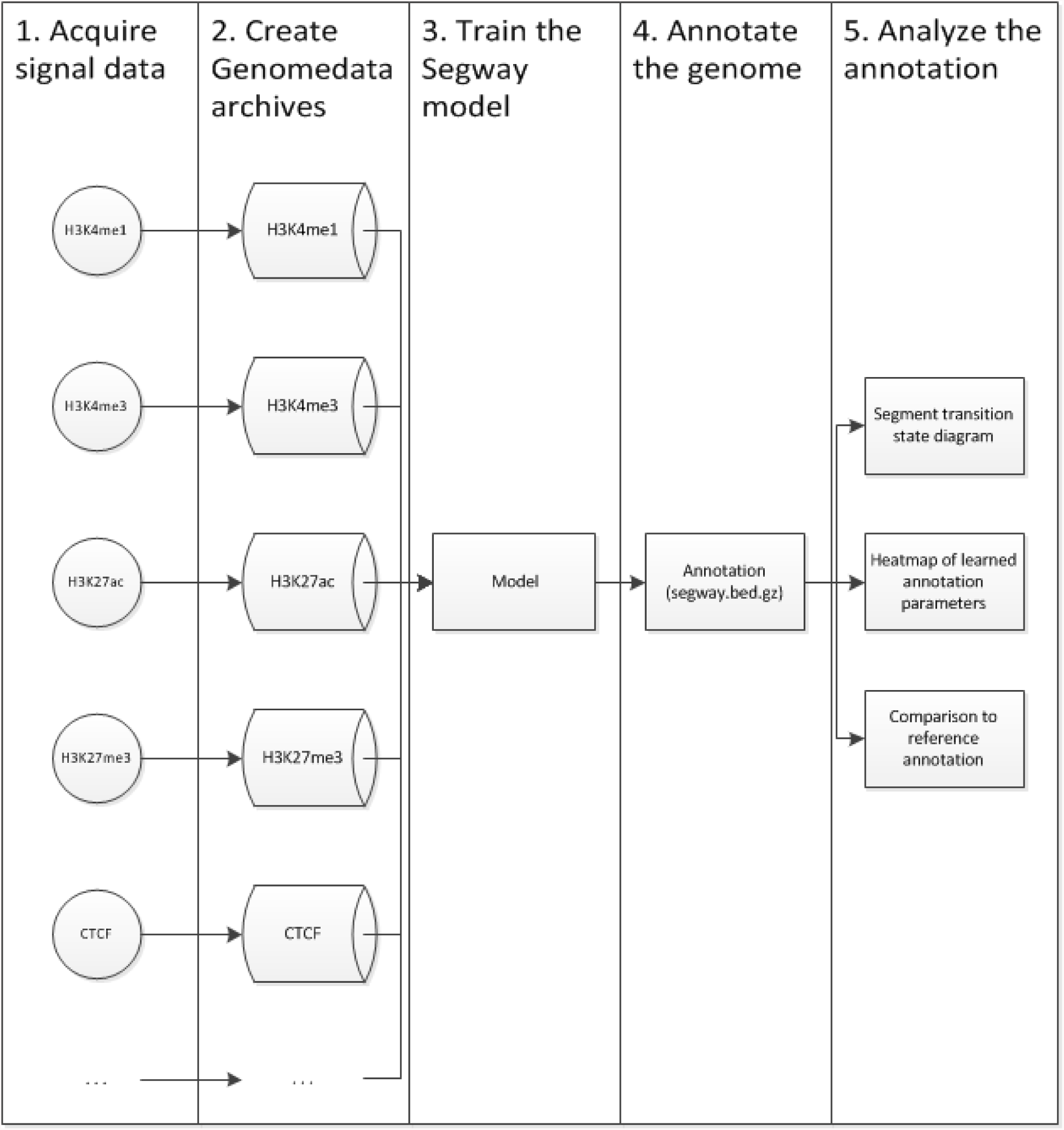
Segway workflow for producing and analysing annotations.

In the PROCEDURE section we illustrate how to perform these steps both for any set of data and specifically on five ChIP-seq experiments for a human B-cell lymphoma cell line, DOHH2^22^.

### Existing Segway annotations

Segway has already produced a number of useful segmentations that are freely available for downloading or viewing in a genome browser. Several of these are available from the Segway website (segway.hoffmanlab.org). The Ensembl Regulatory Build^23^ has Segway annotations of chromatin state across 74 cell types. You can view the Regulatory build segmentations both in Ensembl^24^ and in the UCSC Genome Browser^25^.

### Adapting Segway to other tasks

While most published examples of Segway’s use involve semi-automated genome annotation of chromatin state, it is highly adaptable. One can perform semi-automated genome annotation on any kind of genomic signal data. Since you can also supply an arbitrary Graphical Model Toolkit^26^ (GMTK) dynamic Bayesian network (DBN) model, you can also use Segway as a framework for various different inferences on genomic signal data^27^.

### Primary audience

This protocol was designed for bioinformaticians and other biologists who wish to produce genomic annotations automatically. The signal data can come from public resources, such as ENCODE, or from your own experiments. You should have Linux experience.

## MATERIALS

### EQUIPMENT AND SOFTWARE

- Linux server or workstation
- At least 100 GB of free disk space
- At least 4 GB of memory
- Internet connection
- A Bash shell
- (Recommended) Debian 8, Red Hat Enterprise Linux 7, or CentOS 7
- (Recommended) Cluster system running Grid Engine, IBM Platform Load Sharing Facility (LSF), Portable Batch System (PBS), or Torque

#### Required data

- ChIP-seq read alignments in Binary Alignment/Map (BAM) format (Box 2) or ChIP-seq sequence data tracks from a public source such as ENCODE (Box 3).

## PROCEDURE

### Load signal into Genomedata archives • Timing < 4.5 h

1| Install the prerequisite software in Box 1. This procedure requires that you or your system administrator installed the software and that you have access and permissions to use it.

2| Download (or generate) a *chromosome sizes* file for your genome. This is a tab-delimited file listing all the chromosomes or scaffolds in your genome assembly with lengths measured in base pairs (Figure 2).

**Figure 2:**
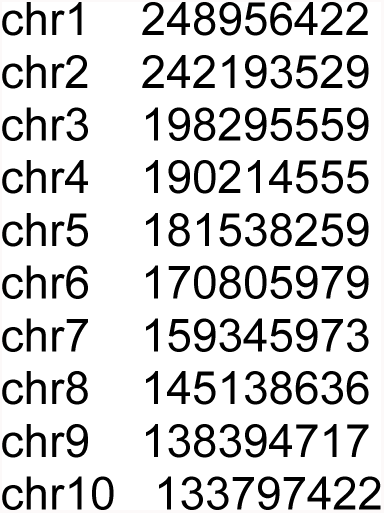
The first 10 lines of the GRCh38 chromosome sizes file used by the ENCODE project.

In this protocol, you will use the Genome Reference Consortium^28^ human genome assembly version GRCh38/hg38, without alternate loci. This assembly also includes autosomal chromosomes 1 to 22, sex chromosomes X and Y, unplaced and unlocalized scaffolds, the mitochondrial chromosome, and the Epstein–Barr virus (“chrEBV”). For this assembly, the ENCODE Project^29^ created a chromosome sizes file (in accession ENCSR425FOI, https://www.encodeproject.org/files/GRCh38_EBV.chrom.sizes/@@download/GRCh38_EBV.chrom.sizes.tsv). To download this file, execute:

~~~
wget https://www.encodeproject.org/files/GRCh38_EBV.chrom.sizes/@@download/GRCh38_EBV.chrom.sizes.tsv -O GRCh38_EBV.chrom.sizes.tsv
~~~

##### □ Critical Step

The genome assembly version used to align all of your signal data must match each other and the sizes file. Always check that assembly versions match and never assume. Failing to ensure consistency will yield nonsensical results from Segway, or any other genome analysis software. There is no guarantee that Genomedata will warn of data in unexpected positions.

3| Download DOHH2 cell line signal data tracks CTCF, H3K4me1, H3K4me3, H3K27ac, H3K27me3 as outlined in Box 3 or generate the signal files from raw reads as outlined in Box 2.

4| Convert any signal files in bigWig format to bedGraph with bigWigToBedGraph. For example, to convert ENCFF884IIL.bigWig to bedGraph format, execute:

~~~
bigWigToBedGraph ENCFF884IIL.bigWig ENCFF884IIL.bedGraph
~~~

5| Filter the signal files for uniquely mappable regions in order to remove signal from repeat regions. The Umap project provides lists of uniquely mappable regions for various assemblies and different genomic dataset read lengths. Download the list of uniquely mappable regions from the Umap^30^ project for your genome and corresponding to the closest read length of your data, (use the smaller read length in case of ties), and convert it to bedGraph. Execute:

~~~
wget
https://www.pmgenomics.ca/hoffmanlab/proj/bismap/raw/hg38/k36.Umap.MultiTrackMappability.wg.gz
bigWigToBedGraph k36.Umap.MultiTrackMappability.bw k36.Umap.MultiTrackMappability.bedGraph
~~~

This Umap file contains a multi-read mappability score, which is the probability that a randomly selected read of a fixed length in a given region is uniquely mappable. The higher the multi-read mappability score, the more confident we are about our observed ChIP-seq signal.

Filter the Umap file for uniquely mappable regions with a multi-read mappability score greater or equal to 0.75. This score is used to ensure that only regions with at least a 75% chance of being correctly mapped are included in the analysis. Execute:

~~~
awk 'BEGIN {FS=OFS=“\t”} {if ($4 >= .75) print $1, $2, $3}'
k36.Umap.MultiTrackMappability.bedGraph | bedtools merge >
k36.Umap.MultiTrackMappability.filtered.bed
~~~

Filter the signal file for uniquely mappable regions using bedtools. For example, to filter ENCFF884IIL.bedGraph, execute:

~~~
bedtools intersect -a ENCFF000BXQ.bedGraph -b
k36.Umap.MultiTrackMappability.filtered.bed >
ENCFF884IIL_from_umap_regions.bedGraph
~~~

##### □ Critical Step

Signal files downloaded from ENCODE (from Box 1), or generated from Box 2 do not distinguish true zero-valued regions from unmappable regions. MACS2^8^ does not distinguish true zero-values from missing data in its output. Segway attempts to accurately and separately model missing data versus zero-valued data. Ideally, your signal files should contain missing data when data is actually missing (by omitting the data) and zero-valued data when there is a known zero value result. If it is impossible to distinguish between zero-valued data and missing data, we recommend setting all missing data to zero. Avoid removing zero-valued data, if possible.

6| Create Genomedata archives containing your reference assembly and signal tracks. Genomedata ^20^ is a format for dense genomic signal data that allows efficient random access. Specifically, create an individual archive for each individual signal track. To create a

Genomedata archive with the GRCh38 human assembly from ENCODE and one of the gathered and filtered signal files from **Step 5**, execute:

~~~
genomedata-load --sizes --sequence GRCh38_EBV.chrom.sizes.tsv --track
H3K4me1=ENCFF884IIL_from_umap_regions.bedGraph DOHH2_H3K4me1.genomedata
~~~

This will result in a directory called DOHH2_H3K4me1.genomedata containing signal and assembly data. The --track option from the previous command allows to assign a name to a track for a given signal file. In this case, it assigns the track name of H3K4me1 to the signal data found in ENCFF884IIL.bedGraph.

7| Create unique named Genomedata archives for the remaining signal files by repeating **Step 6** for each remaining signal file gathered in **Step 3**. If you do this with the suggested datasets, you will have five new Genomedata directories containing the signal data (DOHH2_H3K4me1.genomedata, DOHH2_H3K4me3.genomedata, DOHH2_H3K27me3.genomedata, DOHH2_H3K27ac.genomedata, and DOHH2_CTCF.genomedata).

### Train Segway model from data • Timing < 12 h

8| (Optional) Set a random seed. Segway optimizes model parameters from randomly selected initial values. Usually it is better to let this random selection work unconstrained, but to reproduce the **ANTICIPATED RESULTS** here exactly, you must ensure the same sequence of random numbers. Do this by setting the SEGWAY_RAND_SEED to a positive (32 bit) integer. For example, to replicate **ANTICIPATED RESULTS**, execute:

~~~
export SEGWAY_RAND_SEED=22426492
~~~

**? TROUBLESHOOTING**

9| (Optional) Limit Segway’s processor usage. To do this, set the SEGWAY_NUM_LOCAL_JOBS environment variable to the maximum number of processes you wish Segway to use. This only applies to users who run Segway without a cluster environment such as Grid Engine^31^, and we recommend using such an environment if possible (Box 5). If you need to find out more information about your own processor, execute:

Choose a number no larger than the number of (effective) cores on your machine. This number is the product of the number of processors and each processor’s number of Central Processing Unit (CPU) cores (Figure 3). Smaller values for SEGWAY_NUM_LOCAL_JOBS will result in slower running times and therefore the protocol will take longer to perform. Export an environment variable containing the maximum available number of cores to use for other aspects of the protocol, execute:

~~~
cat /proc/cpuinfo
~~~

**Figure 3:**
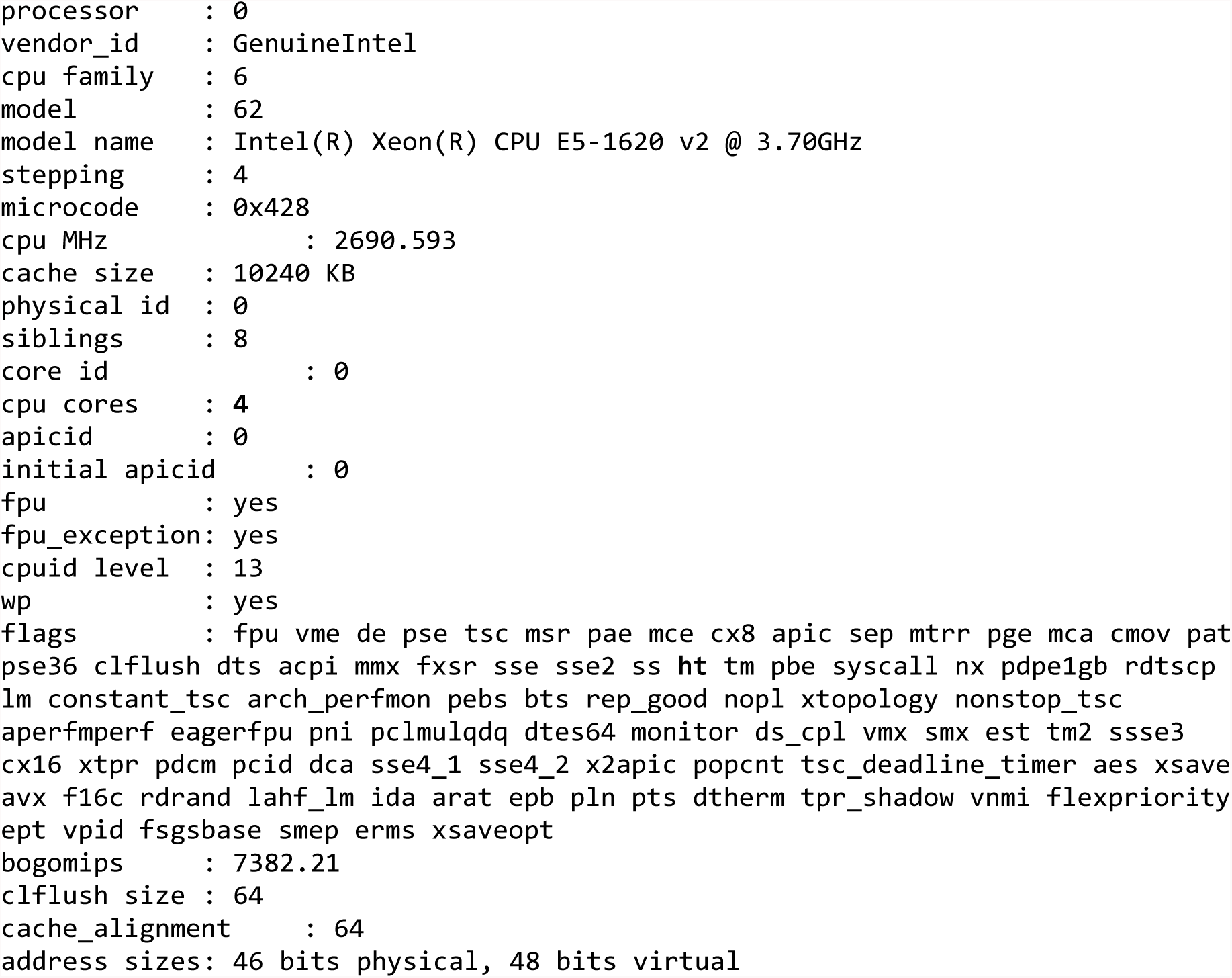
Sample output of “cat /proc/cpuinfo” on a Linux operating system. This particular CPU has two physical cores and hyperthreading (ht), marked in bold above. Hyperthreading allows software to treat one physical core as two effective cores for the operating system. As a result, this machine has four effective cores, marked in bold above.

Choose a number no larger than the number of (effective) cores on your machine. This number is the product of the number of processors and each processor’s number of Central Processing Unit (CPU) cores (Figure 3). Smaller values for SEGWAY_NUM_LOCAL_JOBS will result in slower running times and therefore the protocol will take longer to perform. Export an environment variable containing the maximum available number of cores to use for other aspects of the protocol, execute:

~~~
export NUM_THREADS=$(getconf _NPROCESSORS_ONLN)
~~~

On a cluster, use instead the number of slots allocated to your job. You can also use this variable for SEGWAY_NUM_LOCAL_JOBS by executing:

~~~
SEGWAY_NUM_LOCAL_JOBS=“$NUM_THREADS”
~~~

10| Create regions to exclude from the segmentation. These regions include blacklisted regions for a particular genome or regions you wish to ignore in your analysis. For this protocol, exclude the Epstein–Barr virus (EBV) sequence, because the UCSC Genome Browser does not include the “chrEBV” chromosome.

Create a BED file that excludes the EBV sequence called exclude_coords.bed with the following tab-delimited text:

~~~
chrEBV 0 171823
~~~

11| (Recommended, Optional) Merge a blacklist for your genome assembly into your exclude coordinates. Functional genomic experiments often produce artifact signal in certain regions of the genome. If there is a curated list of these blacklisted regions for the genome and assembly you are using in your annotation, we recommend excluding these regions from your analysis. To merge a blacklist blacklist.bed.gz with the excluded EBV sequence from **Step 10** into a single exclude_coords.bed file, execute:

~~~
zcat -f blacklist.bed.gz ebv_exclude.bed | cut -f 1-3 | sort -k1,1 -k2,2n >
exclude_coords_sorted.bed
bedtools merge -i exclude_coords.sorted.bed > exclude_coords.bed
~~~

12| Train a 10-label Segway model using 1% of your genome. The training process automatically discovers recurring patterns in the signal data you supplied. Training relies on an expectation-maximization^32^ (EM) process that seeks a local maximum likelihood. The likelihood is the probability of generating the given data from the model and its learned parameters. Segway can optimize from multiple sets of initial values simultaneously. Each simultaneous training *instance* results in locally optimized parameters and Segway picks the winner with the best likelihood.

For increased speed, we can train on only a fraction of the genome. Choose a subset to train on using the *minibatch* feature and specifying which fraction of your data you wish to use. The minibatch feature uses a different randomly selected part of the genome in each training round. In Segway we set the number of training rounds with the --max-train-rounds. In most cases, the patterns found after five rounds are quite similar to those after 100. To increase the speed of this example, we will set the maximum number of rounds to 10. Here, we can also speed up training by reducing the resolution of the signal data with the --resolution option.

Create a 10-label model training on 1% of the genome from the archives created from **Steps 6 and 7**, excluding regions from **Steps 10 and 11**, at 10 base pair resolution, using 10 simultaneous training instances by executing:

~~~
segway --resolution=10 --num-instances=10 --minibatch-fraction=0.01 --num-labels=10 --max-train-rounds=10 --exclude-coords=exclude_coords.bed train
DOHH2_H3K4me1.genomedata DOHH2_H3K4me3.genomedata DOHH2_H3K27ac.genomedata K562_H3K27me3.genomedata
DOHH2_CTCF.genomedata train_results
~~~

Segway prints a log of genomic regions it trains on and individual training jobs run on your cluster or in local mode (Figure 4).

**Figure 4:**
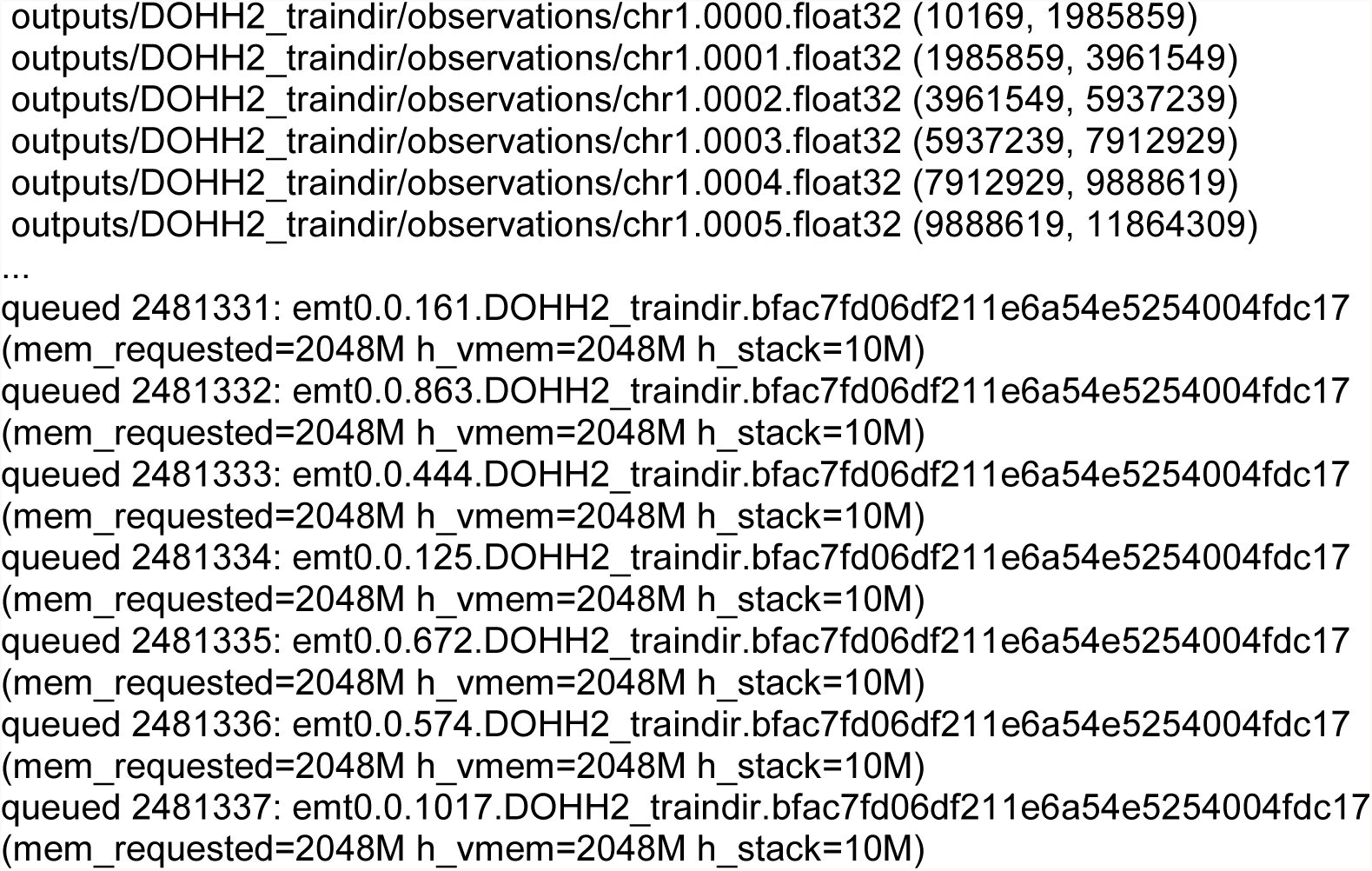
Sample of the expected output from Segway training. The first lines are windows saved for consideration while training the Segway model. The last lines are the EM training (EMT) jobs submitted to a cluster system. Segway determines individual EMT job names using numbers of training instance, EMT round, and window.

**Figure 5:**
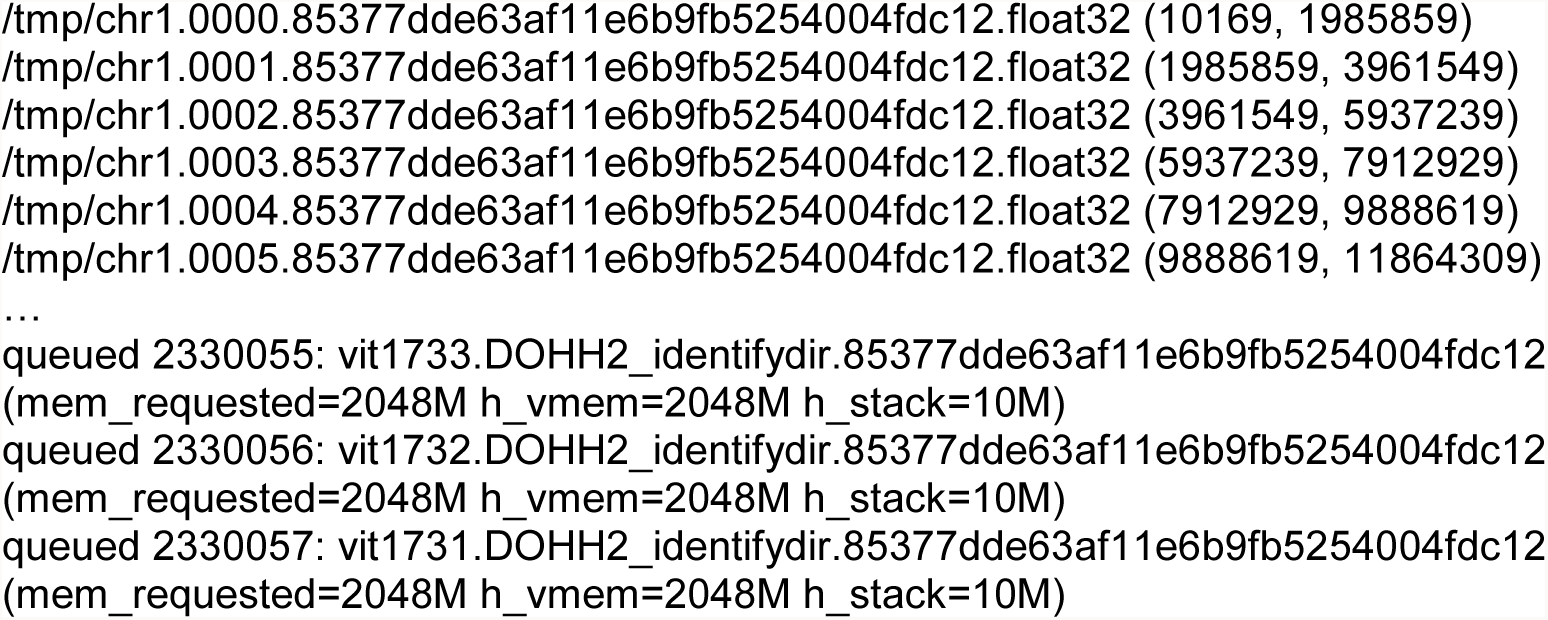
Sample of expected output from the Segway identify task. The first lines are indexed genomic windows saved for subsequent annotation. The last lines are Viterbi jobs to run on a cluster system. Segway numbers the Viterbi job names based on the previously indexed windows.

**? TROUBLESHOOTING**

### Annotate the genome using the trained model • Timing < 12 mins

13| Annotate the genome using the trained model from **Step 12**. The train_results directory contains the final model and trained parameters. To annotate the whole genome from our previously trained model, excluding EBV and blacklist regions from **Steps 10 and 11**, execute:

~~~
segway --exclude-coords="exclude_coords.bed" --
bigBed=identify_results/segway.layered.bb identify
K562_H3K4me1.genomedata K562_H3K4me3.genomedata K562_H3K27.genomedata K562_H3K27m3.genomedata
K562_CTCF.genomedata train_results annotate_results
~~~

Segway prints a log of genomic regions it will annotate and individual identification jobs run on your cluster or in local mode.

Segway writes its annotation to a BED file inside the “identify” directory (annotate_results), named segway.bed.gz. This is a tab-delimited file describing the chromosome regions and their corresponding label number (Figure 6).

**Figure 6:**
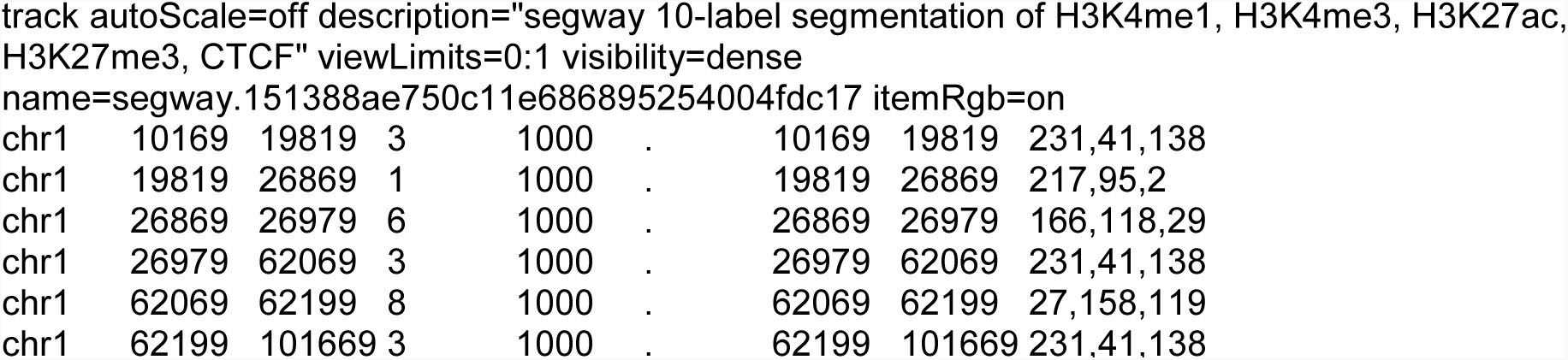
Sample of the resulting segmentation produced by Segway. After a BED file header line, each line contains information on the chromosome, region, and label assigned to each region.

### Analyze the annotation using Segtools • Timing < 30 mins

Segtools ^33^ (segtools.hoffmanlab.org) is a collection of command-line tools that enables exploratory data analysis of genome segmentations, such as the output of Segway. Each tool provides distinct information such as the distribution of segment lengths. Combining Segtools plots and analysis enables you to assign a biological meaning to each annotation label.

14| Plot the emission parameters learned during the training task performed in the **Step 12**.

~~~
segtools-gmtk-parameters train_results/params/params.params
~~~

This creates the gmtk-parameters directory that contains a heatmap (gmtk_parameters.stats.png) showing the learned parameters per label-track pairs. You can run this command directly after the training task.

**? TROUBLESHOOTING**

15| Calculate and plot the length distribution of segments in each label, and the genomic fraction covered by each label using segtools-length-distribution:

~~~
segtools-length-distribution annotate_results/segway.bed.gz
~~~

This creates the length_distribution directory that contains summary statistics in tab-delimited format (length_distribution.tab and segment_sizes.tab) and two plots. The first plot (length_distribution.png) shows the distribution of segment lengths for each label.

The second plot (segment_sizes.png) shows the fraction of total segments for each label and the fraction of genomic bases covered by each label.

16| Calculate the enrichment of each segment label over a gene annotation using segtools-aggregation. Download a gene annotation for your given assembly. In this example, download the GRCh38/hg38 human gene annotation in Gene Transfer Format (GTF) from GENCODE^34^:

~~~
wget
ftp://ftp.sanger.ac.uk/pub/gencode/Gencode_human/release_25/gencode.v25.annotation.gtf.gz
~~~

Calculate and plot the aggregation:

~~~
segtools-aggregation --mode=gene --normalize --outdir aggregate_gene
annotate_results/segway.bed.gz gencode.v25.annotation.gtf.gz
~~~

This command runs segtools-aggregation in “gene” mode and creates two figures showing the enrichment of a segmentation over an idealized transcriptional (aggregate_gene/feature_aggregation.splicing.png) and translational (aggregate_gene/feature_aggregation.translation.png) gene model.

### Visualize signal data and segmentation on the UCSC Genome Browser • Timing < 2 h

The UCSC Genome Browser can load signal tracks and segmentations from a track hub ^35^ —a collection of genome annotation files on a web server. To visualize our segmentation and signal data, make a track hub and visualize it on the UCSC Genome Browser^25^.

17| Create the directory hierarchy for the track hub:

~~~
mkdir -p trackhub/hg38
~~~

This creates the main directory trackhub that will contain all the information necessary for the

UCSC Genome Browser to locate your data. The hg38 subdirectory will contain your Umap-filtered signal tracks and segmentations generated based on the GRCh38/hg38 genome assembly.

18| Create a hub.txt file in the trackhub directory and add the following lines to the beginning of the file:

~~~
hub DOHH2_ChIP-seq
shortLabel DOHH2 ChIP-seq
longLabel Segway annotation of DOHH2 ChIP-seq data
genomesFile genomes.txt
email your.email@example.com
~~~

This file describes the general properties of your track hub where the first word of each line is the name of the property and the rest of the line is the value assigned to it.

19| Create a genomes.txt file in the trackhub directory file describing the genome assembly and the path to the track property file trackDb.txt for that genome assembly:

~~~
genome hg38
trackDb hg38/trackDb.txt
~~~

20| Convert all filtered signal files to the bigWig^36^ format. Track hubs can only load signal data that are in binary index formats, such as bigWig. To convert the previously filtered signal data (in **Step 5**) in bedGraph format to bigWig, use bedGraphToBigWig. This tool requires a sorted input bedGraph file.

Here is an example showing how to convert a bedGraph file to bigWig:

~~~
sort -k1,1 -k2,2n DOHH2_H3K4me1.filtered.bedGraph >
DOHH2_H3K4me1_sorted.filtered.bedGraph
bedGraphToBigWig DOHH2_H3K4me1_sorted.filtered.bedGraph
GRCh38_EBV.chrom.sizes.tsv trackhub/hg38/DOHH2_HK4me1.bigWig
~~~

The first command sorts the signal file DOHH2_H3K4me1.filtered.bedGraph by chromosome and start position, and writes the output to DOHH2_H3K4me1_sorted.filtered.bedGraph. This step fulfills the requirement that the input signal and chromosome sizes files both be sorted in the same manner. GRCh38_EBV.chrom.sizes.tsv is the chromosome sizes file downloaded in the **Steps 1–3** and trackhub/hg38/DOHH2_H3K4me1.bigWig is the name of the output file converted to bigWig. Here we specifically store the output file in the trackhub/hg38 directory created above.

Note that you should execute the same procedure for each filtered bedGraph signal file, replacing DOHH2_H3K4me1 by the signal filename in the example above.

21| Move the layered segmentation generated in **Step 13** to the trackhub/hg38 directory.

Execute:

~~~
mv identify_results/segway.layered.bb trackhub/hg38/
~~~

22| Create trackhub/hg38/trackDb.txt which describes how to display the tracks. Set the following parameters for an optimal view of segmentation tracks.

~~~
track DOHH2_Segway
type bigBed 12
bigDataUrl segway.layered.bb
shortLabel DOHH2_segmentation
longLabel segmentation of DOHH2 cell line from ChIP-seq data
itemRgb on
visibility pack
~~~

Set the following parameters for the signal tracks.

~~~
track DOHH2_H3K4me1
type bigWig
bigDataUrl DOHH2_H3K4me1.bigWig
shortLabel DOHH2_H3K4me1
longLabel ChIP-seq signal in DOHH2
visibility full
maxHeightPixels 100:60:8
viewLimits 0:100
track DOHH2_H3K4me3
type bigWig
bigDataUrl DOHH2_H3K4me3.bigWig
shortLabel DOHH2_H3K4me3
longLabel ChIP-seq signal in DOHH2
visibility full
maxHeightPixels 100:60:8
viewLimits 0:100
track DOHH2_H3K27me3
type bigWig
bigDataUrl DOHH2_H3K27me3.bigWig
shortLabel DOHH2_H3K27me3
longLabel ChIP-seq signal in DOHH2
visibility full
maxHeightPixels 100:60:8
viewLimits 0:100
track DOHH2_H3K27ac
type bigWig
bigDataUrl DOHH2_H3K27ac.bigWig
shortLabel DOHH2_H3K27ac
longLabel ChIP-seq signal in DOHH2
visibility full
maxHeightPixels 100:60:8
viewLimits 0:100
track DOHH2_CTCF
type bigWig
bigDataUrl DOHH2_CTCF.bigWig
shortLabel DOHH2_CTCF
longLabel ChIP-seq signal in DOHH2
visibility full
maxHeightPixels 100:60:8
viewLimits 0:100
~~~

The UCSC Genome Browser provides many other options as described on its website (genome.ucsc.edu/goldenpath/help/trackDb/trackDbHub.html#commonSettings).

23| Upload the track hub directory to a public web space. For example, to copy changes to a remote server named yourserver with username yourname, execute:

~~~
rsync -a trackhub yourname@yourserver:/your/publicly/available/space/trackhub
~~~

The -a option specifies rsync’s archive mode, which preserves all file attributes, recursively copying files and directories.

24| Visit the UCSC Genome Browser (https://genome.ucsc.edu) and load your track hub. To do so, select “My data” > “Track hubs” from the top menu and add the direct link to your hub.txt file in the “URL” field. Push the “Add Hub” button. This will allow you to visualize your segmentation as well as your signal tracks, if you included them in your track hub.

**? TROUBLESHOOTING**

## ? TROUBLESHOOTING

Table 1 contains troubleshooting recommendations.

**Table 1.** Troubleshooting Table

| Step | Problem | Possible Reason | Solution |
| --- | --- | --- | --- |
| 8 | Segway produces identical results each time it is run | Accidentally have the random seed for Segway set | Remove the Segway random seed with the command `unset SEGWAY_RAND_SEED` |
| 8 | Segway produces different results each time it is run | No random seed has been set or not enough instances have been run | If you're attempting to reproduce results consistently, consider using a random seed as described in **Step 8**. If you're not using a seed but getting significantly different results every time Segway is run, increasing your `--num-instances` will allow Segway to create a more consistent trained model since it gains a larger trained sample size |
| 12 | Need to recover from a crashed Segway process | This could be a cluster issue or configuration issue. Check the error log output by Segway for the reason your training/identify run crashed for details | To recover your Segway run, rename your old (train or identify) results directory to a new name which Segway can use to recover from. Specify the new directory name with the `--recover` option. Recovery only works with identical parameters |
| 12 | Not enough disk space or temporary directory issues while or after running | Segway creates temporary observation files as part of its process. | Set the `TMPDIR` environment variable to a directory where you have write |
|  | Segway |  | access and sufficient disk space |
| 14 | Segtools does not produce PNG images | To create PNG images, R requires an X11 display which will not be present on a headless node | Copy the necessary files to a machine that has Segtools and an X11 display available and run Segtools there. Alternatively Install the R package, Cairo, on your machine. This may require a number of additional dependencies to be installed |
| 24 | Track data from the uploaded Track hub is not displaying correctly on the UCSC Genome Browser | The server hosting the track data does not support byte serving or does not advertise it with an "Accept-Ranges" HTTP response header | Contact the hosting administrator to enable byte serving for the track hub track datasets or find an alternative web host that supports byte serving |
| Box 1 | I have Python 2.7 but my system reports that pip is not installed | Pip does not automatically come bundled with some distributions of Python 2.7, or older versions of Python | If possible, ask your administrator to install the necessary python package containing pip. If running Python 2.7.9 or later, it is possible to install pip through the ensurepip module with 'python -m ensurepip --upgrade'. For all other cases refer to the pip documentation |
| Box 1 | There is a Unicode-related error when attempting to install numpy with pip | Older versions of pip cannot install numpy correctly due to a unicode related bug | Upgrade your pip version using 'pip install --upgrade pip' |
| Box 1 | The computer to run Segway on doesn't have an internet | Some cluster systems have strict policies regarding | On a machine with an internet connection download Segway |
|  | connection | internet connectivity | and all dependencies without installation by executing 'pip install --download segway_packages segway'. Copy the segway_packages directory to your target machine without an internet connection. To install Segway from the directory of python packages, execute: 'pip install --no-index --find-links segway_packages segway' |
| Box 2 | MACS2 produces a segfault error, for example, after attempting to 'Call peaks for each chromosome' | MACS2 will not create the necessary output directories for you | Create the directory manually using 'mkdir' and run your MACS2 command again |
| Box 2 | SPP installation fails with the error: 'configure: error: cannot find Boost headers version >= 1.41.0' | SPP is unable to find your Boost C++ installation | Ensure that you have properly set your BOOST_ROOT environment variable to include your Boost C++ installation. For example, if your Boost installation is located in directory sampledir, execute: 'export BOOST_ROOT=sample dir'. |
| Box 4 | ChIPQCreport reports X11 errors | To create images, ChIPQCreport requires an X11 display which will not be present on a headless node | Copy the necessary files to a machine that has Segtools and an X11 display available and run ChIPQCreport there. Alternatively, on a headless node install<br>the R package, Cairo, on your machine. This may require a number of additional dependencies to be installed. Place the necessary commands in a R script file, x.R, and run it via<br>`xvfb-run -s "-screen 0 1600x1200x24+32" Rscript x.R`<br>It may be necessary to also set `options(bitmapType='cairo')` in your R script file |

### • Timing

The entire protocol takes < 29 h, with approximately 1.5 h of configuration and entering commands and approximately 27.5 h of computation. We took the timings for this protocol from a Grid Engine cluster system where we submitted each job to a machine running at 2.6 GHz with 20 MB of cache and eight effective CPU cores.

Step 1, installing the prerequisite software as administrator: < 30 min. Without administrative privileges it takes more time, is largely dependent on the experience of the user, and can range from 30 min to <2 h.

Steps 2–3, downloading data from ENCODE: < 1.5 h. This step largely depends on the speed of the internet connection used to download the datasets. We downloaded the datasets at 3 MB/s– 4 MB/s.

Steps 4–6, filtering data for uniquely mappable regions: 30 min

Step 7, creating the Genomedata archives: 2 h

Steps 8–12, prepare and train Segway with a 10-label model: 12 h

Step 13, annotate the genome with the trained Segway model: 12 h

Steps 14–16, analyzing the annotation with Segtools: 30 min

Steps 17–24, create and upload a trackhub of the annotation: 2 h. The timing on these steps depends on the speed of the internet connection used to upload the datasets.

For smaller datasets not in this protocol, such as K562 GRCh37/hg19 with the same ChIP-seq targets from ENCODE, the computation time is substantially reduced from approximately 24 h to 5 h. The speed of the protocol largely depends on the availability of processors Segway can submit jobs to and the speed of processors themselves. Any bottlenecks on a cluster system or running Segway on a limited number of processors will substantially increase the protocol length.

## ANTICIPATED RESULTS

Segway produces an annotation for a given cell type. We illustrate the results of Segway’s annotation on DOHH2 from **Step 13** by exploring the output of Segtools produced in the **Steps 14–16** and visualizing the segmentation on the UCSC Genome Browser using the track hub produced in **Steps 17–24**.

### Exploring the parameters learned during the training task

The command described in **Step 14** creates the gmtk- parameters/gmtk_parameters.stats.png file showing the Gaussian parameters learned by Segway during the training task described in **Step 12**. The file contains a heatmap (Figure 7) with the data tracks in rows and the labels in columns. For each track-label combination, Segway learns a probability distribution over track values given a label. By default, it uses a Gaussian, or normal distribution, for this probability distribution. The colors on the heatmap represent row-normalized Gaussian means, where dark blue indicates a low mean and dark red indicates a high mean. The sizes of the black rectangles represent the variance parameter of the Gaussian, where larger rectangles indicate a higher variance. For example, label 1 associates with high values for H3K4me3, H3K27ac, and H3K4me1 tracks, shown in red. Low values occur for CTCF and H3K27me3, shown in blue. This observation allows us to hypothesize that label 1 is associated with active genes’ transcription start sites.

**Figure 7:**
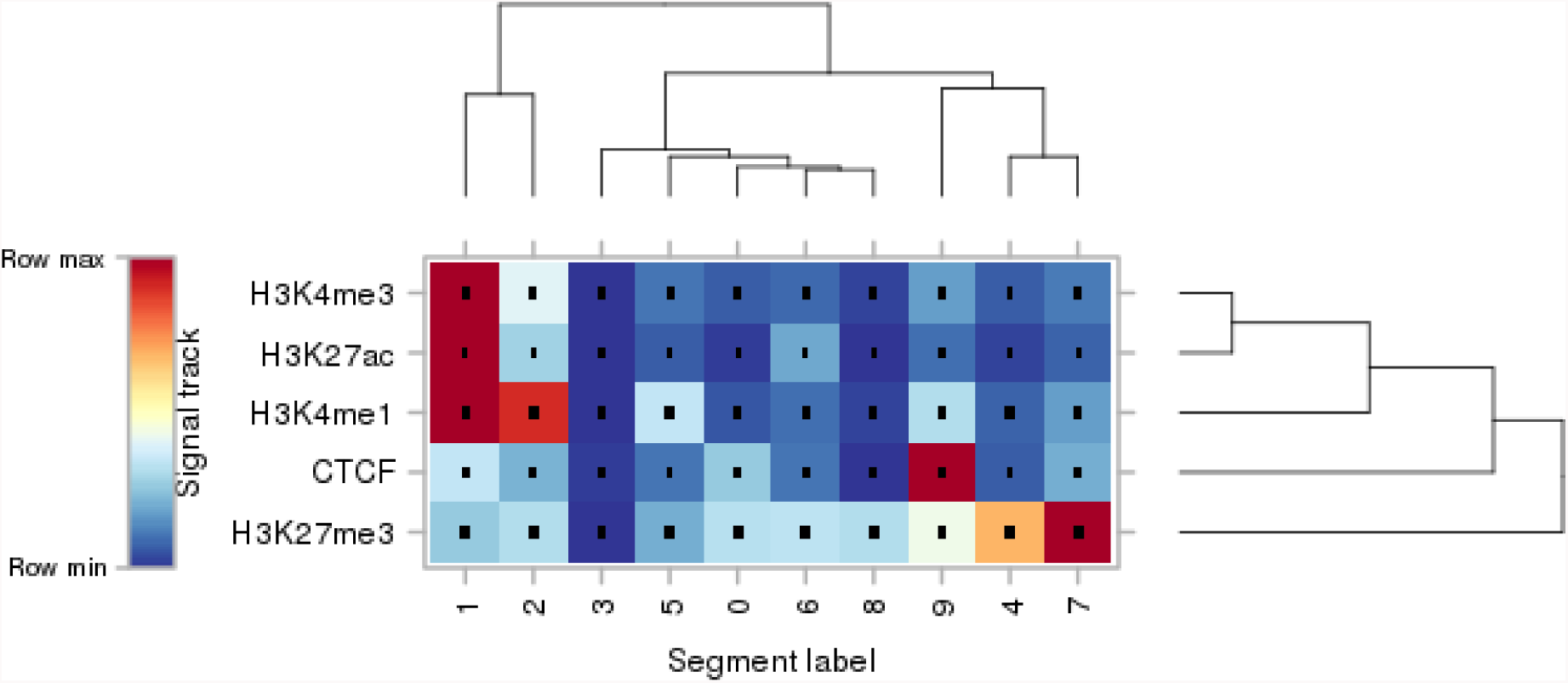
Gaussian emission parameters learned by training a 10-label model on 5-signal dataset.

### Exploring the segment length distribution

The command described in **Step 15** creates the length_distribution directory. This directory contains summary statistics in tab-delimited format (length_distribution.tab and segment_sizes.tab). Segtools uses these statistics to generate summary plots. The first plot (length_distribution.png) (Figure 8) shows the distribution of segment lengths for each label.

**Figure 8:**
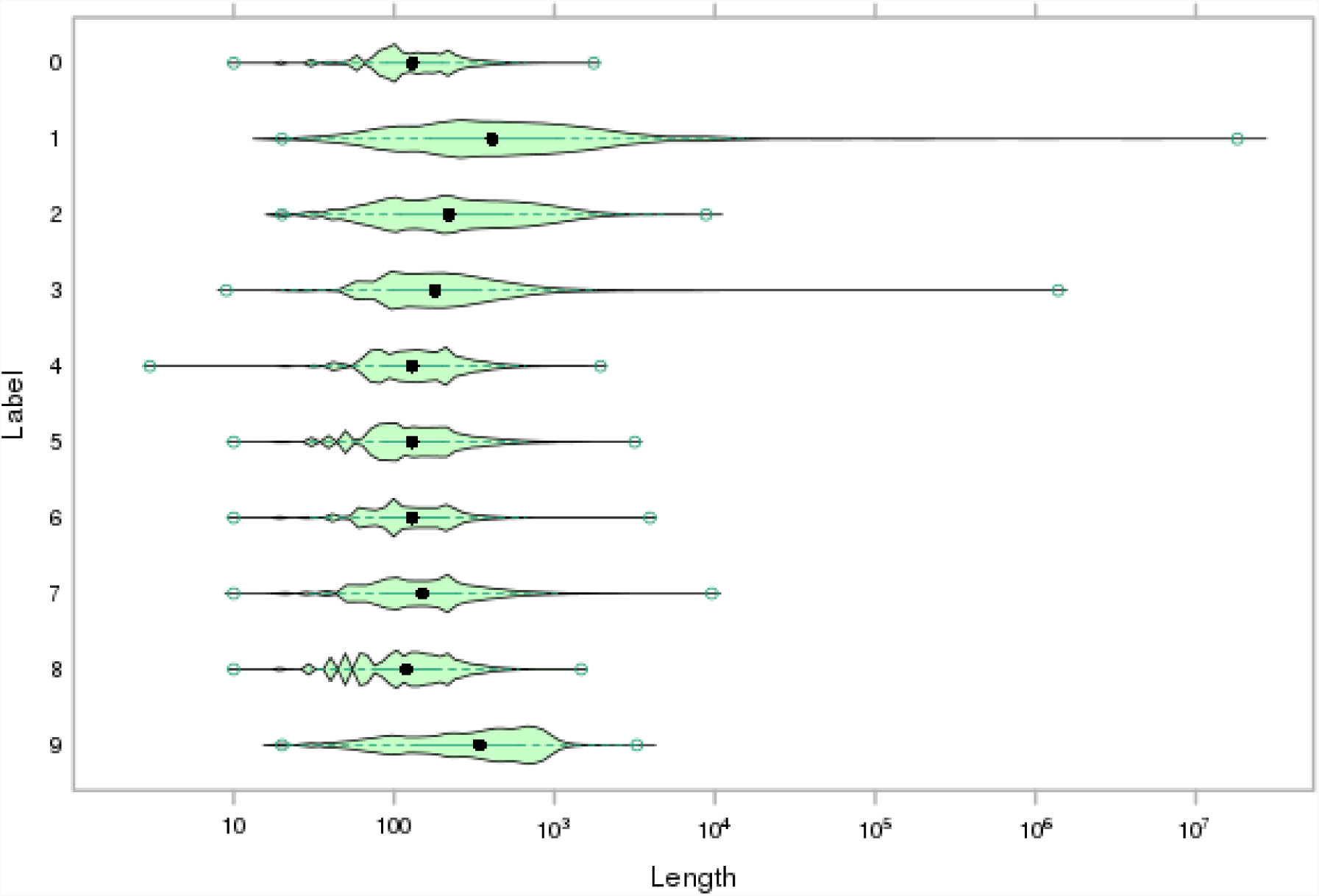
Segment length distribution per label.

The second plot (segment_sizes.png) (Figure 9) shows the fraction of total segments for each label and the fraction of genomic bases covered by each label. Some segments cover very large regions. In particular, Label 1 has some segments with extremely large length. Since the segmentation included some large assembly gaps, Segway picked the default highest average mean label. This is a result of the protocol using a genome sizes file for describing the reference genome used in the analysis and not a more precise description of the genome such as an Assembly Golden Path^37^ (AGP) file where such regions would not be considered for training or annotation.

**Figure 9:**
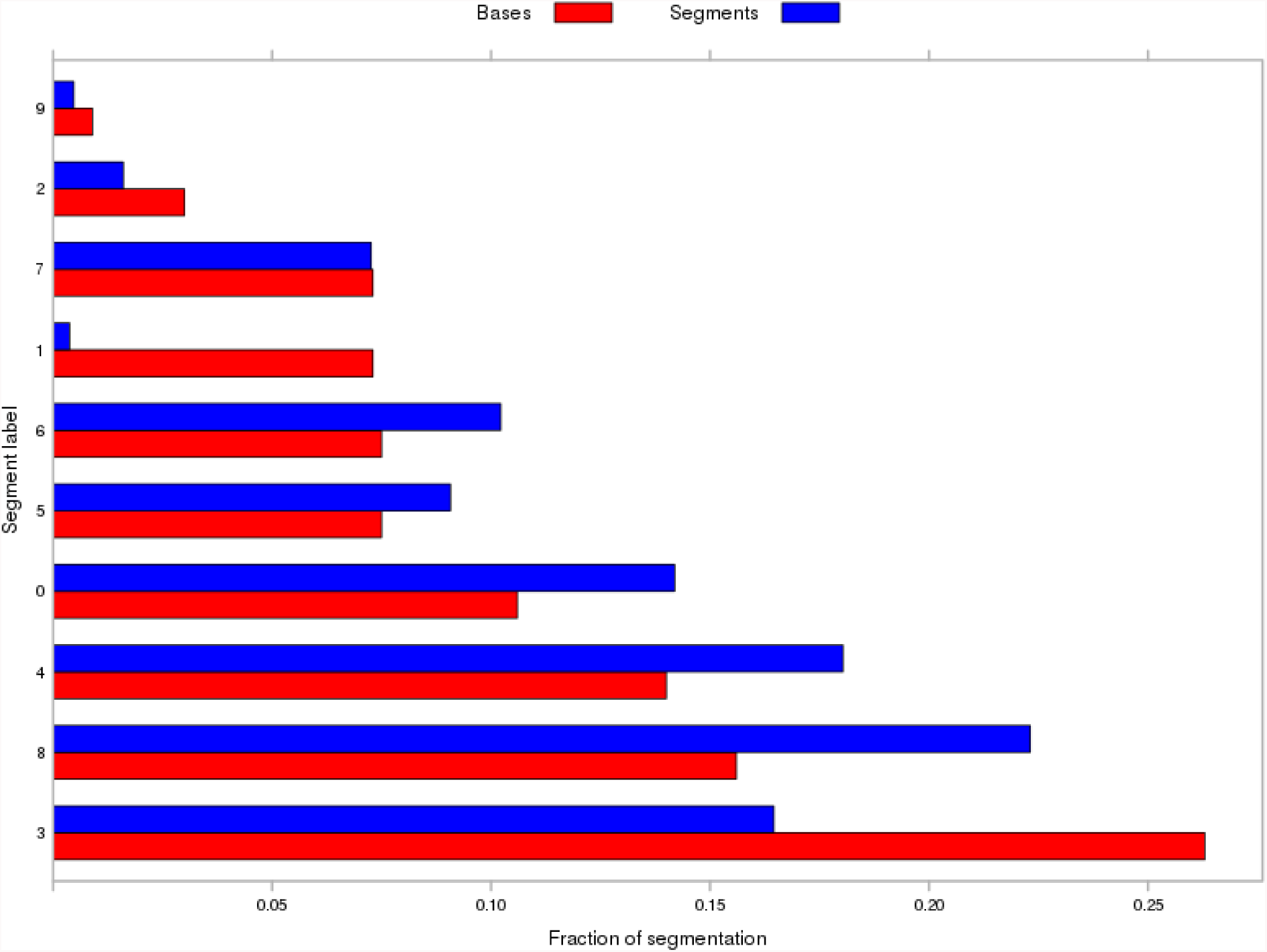
Segment sizes and segment count per labels. In blue: the fraction of the annotation covered for each label by number of segments. In red: the fraction of the segmentation covered by bases for each label.

### Exploring the segment enrichment against gene annotation

The command described in **Step 16** generates the file aggregate_gene/feature_aggregation.splicing.png, which summarizes the occurrence of each segmentation label (y-axis) relative to an idealized transcriptional gene model separated into 8 components (x-axis). For example, one can see the enrichment of label 1, in red, over components generally found around the 5′ end of a gene (Figure 10).

**Figure 10:**
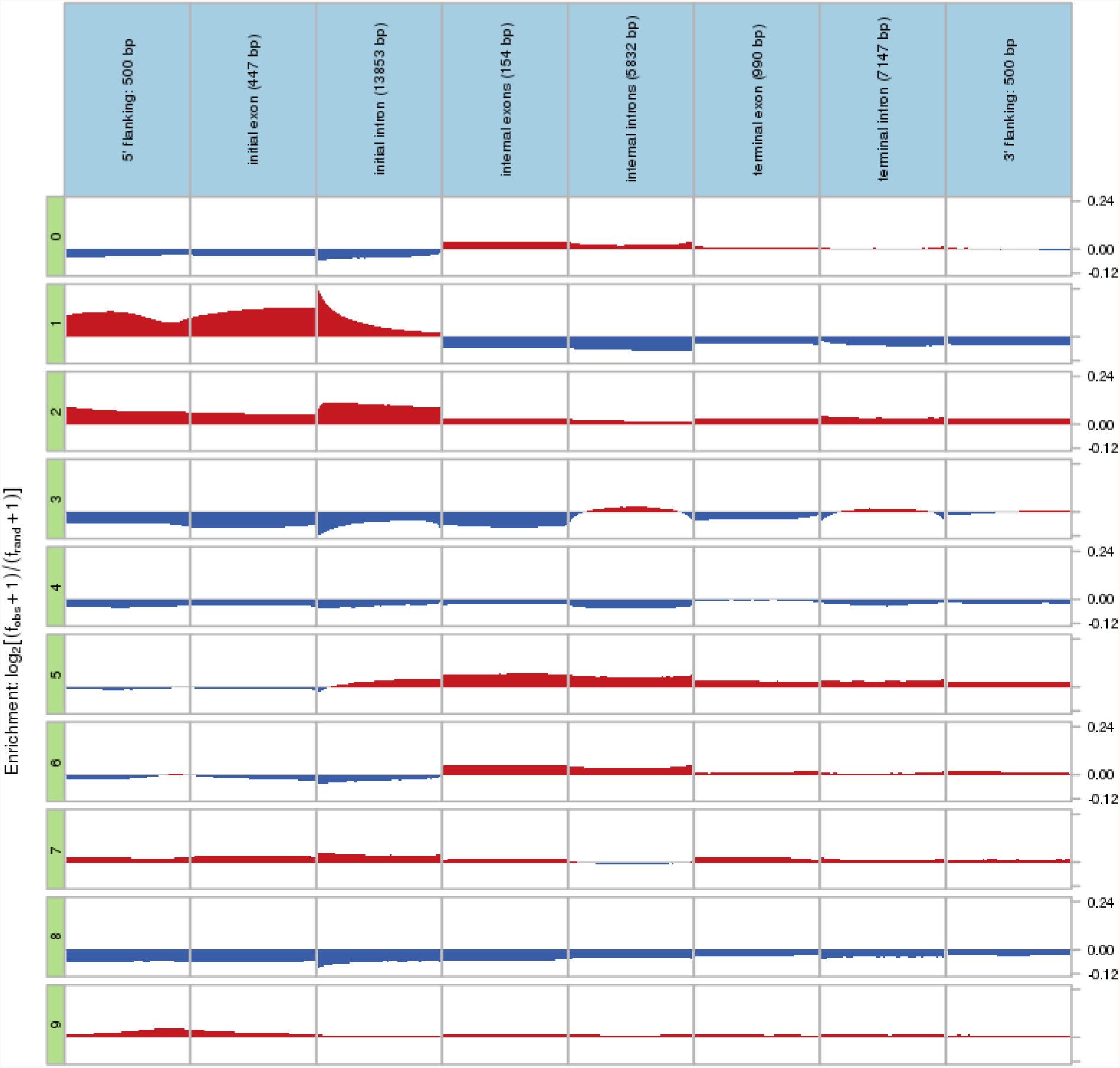
Segment labels’ enrichment relative to an idealized gene model derived from GENCODE 25. Color indicates enrichment (red) or depletion (blue).

### Visualize Segway results on a genome browser

Figure 11 illustrates the track hub generated in **Steps 17–24** loaded on the UCSC Genome Browser. The first track shows the GENCODE annotation of the *CDK1* locus. The subsequent tracks display the signal values from the five bigWig files generated in **Step 20**. Finally, the last track is the segmentation generated with Segway. In this example, labels 1 and 2 are present around the 5′ end of the *CDK1* gene. Labels 5 and 6 cover the middle and end of the gene.

**Figure 11:**
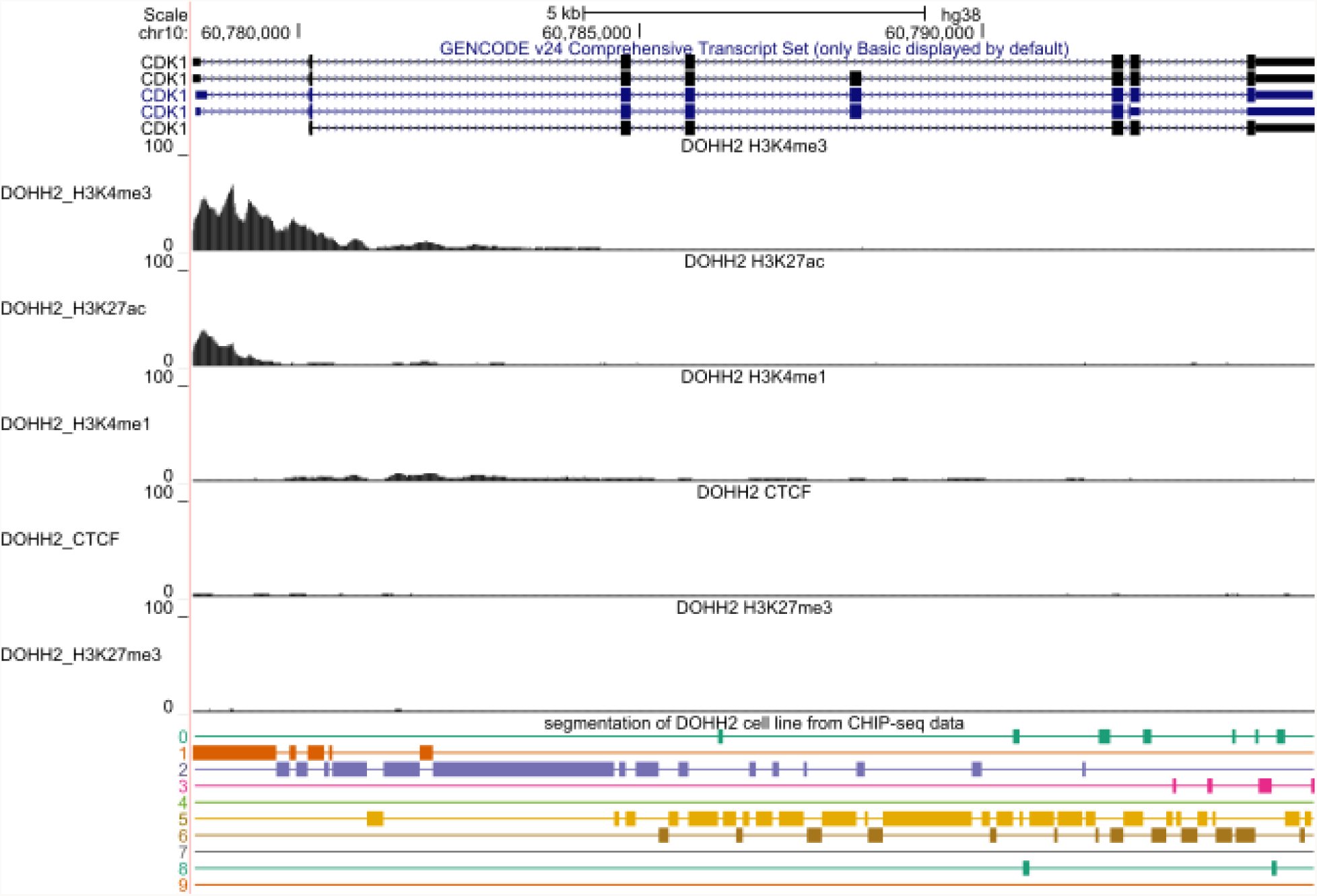
Example of signal tracks and segmentations on the UCSC Genome Browser for the *CDK1* locus.

These observations are consistent with the enrichments shown in Figure 10.

## Acknowledgments

We thank Carl Virtanen and Zhibin Lu (University Health Network High Performance Computing Centre and Bioinformatics Core) for technical assistance and J. Seth Strattan for assistance with the signal datasets used in this protocol. We thank Thomas Carroll for providing support for ChIPQC and for expediting work on ChIPQC’s GRCh38 annotation.

This work was supported by the Canadian Cancer Society (703827 to M.M.H.), the Ontario Institute of Cancer Research (OICR) through funding provided by the Government of Ontario (CSC-FR-UHN to John E. Dick), the Ontario Ministry of Research, Innovation and Science (ER15-11-233 to M.M.H.), the Natural Sciences and Engineering Research Council of Canada (RGPIN-2015-03948 to M.M.H. and an Alexander Graham Bell Canada Graduate Scholarship to C.V.), the Ontario Ministry of Training, Colleges and Universities (Ontario Graduate Scholarship to C.V.), the University of Toronto Medicine by Design (C1TPA-2016-04 to M.M.H. and C1TPA-2016-01 to M.M.H.), the McLaughlin Centre (MC-2015-16 to M.M.H.), the National Institutes of Health (GM119550 to J.R.H.), the American Cancer Society (RSG-13-216-01-DMC to J.R.H.), and the Princess Margaret Cancer Foundation.

## Author Contributions

E.G.R. wrote the introduction steps 1–4, 6–13 and Boxes 1 and 5.

M.M. wrote steps 5 and 14–24 of the procedure and the anticipated results.

E.G.R. and R.C. tested the protocol and improved the software.

M.K. and D.C. provided additional troubleshooting and testing.

A.K., R.A, R.C, and J.R.H wrote Box 2.

M.K. wrote Box 3.

C.V. wrote Box 4

R.A. and J.H. wrote Box 6.

E.G.R. and M.M.H. designed and coordinated the production of the overall protocol.

## Footnotes

**Competing Financial Interests** The authors declare that they have no competing financial interests.

## Boxes

#### Box 1 Installing Prerequisite Software

Segway is only supported on Linux.

There are two distinct methods to install Segway on your machine.

1. Install Segway as an administrator, for all users on the system. This is the easier method, if you have the administrator privileges.
2. Install Segway without any special privileges, for one user.

For either method you need to install:

- Python (2.7)
- Hierarchical Data Format 5 (HDF5) (1.8.17)
- Graphical Models Toolkit (GMTK) (1.4.2)
- - Segway (1.4.1)
- - R (3.3)
- Segtools (1.11)
- bedtools (2.26.0)
- bigWigToBedGraph
- bedGraphToBigWig (4)
- bedToBigBed (2.7)

##### Installing Segway as administrator

1. Install Python (with pip), HDF5, and R with your system package manager

##### Ubuntu or Debian 8

sudo apt-get install python2.7 libhdf5-serial-dev hdf5-tools r-base bedtools python-pip

##### CentOS 7, Red Hat Enterprise Linux 7, Fedora

sudo yum -y install hdf5 hdf5-devel R BEDTools readline-devel python-devel

For Red Hat Systems we recommend using the existing installed version of Python 2.7.

Upgrading the system Python can break yum. **For Fedora 22+ only:** replace ‘yum’ with

‘dnf’.

##### Install R dependencies

From an interactive R environment (with sudo access):

install.packages(c(“latticeExtra”, “reshape2”), repos='http://cloud.r-project.org/')

##### All Linux Distributions

###### GMTK

~~~
wget http://melodi.ee.washington.edu/downloads/gmtk/gmtk-1.4.4.tar.gz
tar xf gmtk-1.4.4.tar.gz
cd gmtk-1.4.4
./configure
make
make install
~~~

##### Segtools

sudo pip install segtools

##### Segway

On Debian only:

~~~
export LD_LIBRARY_PATH=$LD_LIBRARY_PATH:"/usr/lib/x86_64-linux-gnu/hdf5/serial"
export C_INCLUDE_PATH=$C_INCLUDE_PATH:"/usr/include/hdf5/serial"
sudo pip install segway
~~~

##### Installing Segway without administrator privileges

We highly recommend, when possible, getting an administrator to install the necessary software. Each individual software package may have dependencies not installed on your machine. In case this is not possible, as a backup procedure, we describe how to install some of the required software, assuming your system has the prerequisites of that software.

For each prerequisite not already installed, to install without administrator privileges, execute the following steps:

1. Download the software source code
2. Decompress and extract to a directory, if necessary
3. Configure the software source code to where you want to install the software
4. Build and install the software itself
5. Prepend the installation directory of your software on your PATH environment variable.

As an example, to install for Python 2.7.11 the full set commands to install to ~/.local/python2.7.11 execute:

~~~
wget https://www.python.org/ftp/python/2.7.11/Python-2.7.11.tgz
tar xf <u>Python-2.7.11.tgz</u>
cd Python-2.7.11
./configure --prefix=${HOME}/.local/python2.7.11
make
make install
export PATH="${HOME}/.local/python2.7.11/bin:${PATH}"
~~~

You will probably want to place the export PATH command inside your shell configuration file, such as ~/.bashrc

Run similar commands for installing HDF5 and GMTK into their own directories as well. The download locations are:

GMTK - https://melodi.ee.washington.edu/gmtk/

HDF5 - https://www.hdfgroup.org/downloads/index.html

R - http://cloud.r-project.org/mirrors.html

Bedtools - https://github.com/arq5x/bedtools2/releases

Commands to install each prerequisite follow:

##### GMTK

~~~
wget http://melodi.ee.washington.edu/downloads/gmtk/gmtk-1.4.4.tar.gz
tar xf gmtk-1.4.4.tar.gz
cd gmtk-1.4.4
./configure --prefix=${HOME}/.local/gmtk-1.4.4
make
make install
export PATH="${HOME}/.local/gmtk-1.4.4/bin:${PATH}"
~~~

##### HDF5

~~~
wget http://www.hdfgroup.org/ftp/HDF5/current/src/hdf5-1.8.17.tar.gz
tar xf hdf5-1.8.17.tar.gz
cd hdf5-1.8.17
./configure --prefix=${HOME}/.local/hdf5-1.8.17
make
make install
export PATH="${HOME}/.local/hdf5-1.8.17/bin:${PATH}"
export HDF5_DIR="${HOME}/.local/hdf5-1.8.17"
export LD_LIBRARY_PATH="${HOME}/.local/hdf5-1.8.17/lib:${LD_LIBRARY_PATH}"
~~~

##### R

~~~
wget http://cloud.r-project.org/src/base/R-3/R-3.3.0.tar.gz
tar xf R-3.3.0.tar.gz
cd R-3.3.0
./configure --prefix=${HOME}/.local/R-3.3.0
make
make install
export PATH="${HOME}/.local/R-3.3.0/bin:${PATH}"
~~~

##### R dependencies

From an interactive R environment:

~~~
install.packages(c(“latticeExtra”, “reshape2”), repos='http://cloud.r-project.org/')
~~~

##### Bedtools

~~~
wget https://github.com/arq5x/bedtools2/releases/download/v2.26.0/bedtools-2.26.0.tar.gz
tar xf bedtools-2.26.0.tar.gz
cd bedtools2
export prefix=${HOME}/.local/bedtools-2.26
make install
export PATH="${HOME}/.local/bedtools-2.26/bin:${PATH}"
~~~

##### Segway and Segtools

Execute the following to install Segway and Segtools:

~~~
pip install --user segtools
pip install --user segway
~~~

Python software installed this way with pip will place executables in ~/.local/bin. Prepend this to your PATH environment variable.

##### Installing remaining utilities

The remaining software prerequisites are utilities distributed as standalone binaries. You will use these utilities to convert genomic signal and annotation data between different formats. Here we will install the utilities in your home directory in .local/bin. If the directory doesn’t exist, create the directory with the following command: mkdir -p “${HOME}/.local/bin"

Ensure your PATH environment variable contains the location of your downloaded binaries. To place ~/.local/bin in your PATH, execute:

~~~
export PATH="${HOME}/.local/bin:${PATH}"
~~~

Install the remaining utilities with the following commands:

##### bigWigToBedGraph

~~~
wget http://hgdownload.cse.ucsc.edu/admin/exe/linux.x86_64/bigWigToBedGraph
chmod +x bigWigToBedGraph
mv bigWigToBedGraph “${HOME}/.local/bin"
~~~

##### bedGraphToBigWig

~~~
wget http://hgdownload.cse.ucsc.edu/admin/exe/linux.x86_64/bedGraphToBigWig
chmod +x bedGraphToBigWig
mv bedGraphToBigWig “${HOME}/.local/bin/"
~~~

##### bedToBigBed

~~~
wget http://hgdownload.cse.ucsc.edu/admin/exe/linux.x86_64/bedToBigBed
chmod +x bedToBigBed
mv bedToBigBed "${HOME}/.local/bin"
If any installation step fails, refer to ? TROUBLESHOOTING.
~~~

#### Box 2 Generate ChIP-seq Signal Files Using MACS2

Segway takes its input from signal files—normalized representations of ChIP-seq reads within a genomic region. You can convert aligned ChIP-seq data in BAM format to signal files in bedGraph or bigWig format. This protocol uses bedGraph format.

You should compute predominant fragment lengths for the ChIP-seq data prior to generating the fold-enrichment signal files using the MACS2 software. We compare fragment-lengths to read-lengths in order to provide estimates about the amount of background signal in the ChIP-seq data.

##### Estimate predominant fragment length

The program phantompeakqualtools^38,39,40^ (v1.1) will calculate the predominant insert-size (or fragment length) based on strand cross-correlation analysis. The program package is available at https://github.com/kundajelab/phantompeakqualtools and includes detailed installation instructions. This program runs on a Unix system and requires the following dependencies: R (>=3.1), awk, samtools, boost C++ libraries, R packages: spp (>=1.14), caTools, snow.

To download the phantompeakqualtools program to your local environment, execute:

~~~
wget https://github.com/kundajelab/phantompeakqualtools/archive/master.zip
-O phantompeakqualtools.zip
unzip phantompeakqualtools.zip
~~~

From an interactive R environment (see installation instructions in Box 1), execute the following commands to install the phantompeakqualtools program and its required R package dependencies:

~~~
install.packages(c(“caTools”, “snowfall”), repos='http://cloud.r-project.org/')
source(“http://bioconductor.org/biocLite.R”) biocLite(“Rsamtools”)
install.packages(“phantompeakqualtools-master/spp_1.14.tar.gz”)
~~~

##### Installing Sambamba

The Sambamba^41^ program is used to process BAM files. If your files are in tagAlign format then you do not need to use Sambamba, as shown below. You can download Sambamba from GitHub (https://github.com/lomereiter/sambamba/releases). For example, to install Sambamba on a 64-bit Linux machine, execute

~~~
wget -qO-
https://github.com/lomereiter/sambamba/releases/download/v0.6.3/sambamba_v0_linux.tar.bz2 | tar xj -C $HOME/bin/ --transform ’s/_v.*//'
~~~

##### Generating a tagAlign file

Prior to generating the fragment length estimation, execute the following to generate a tagAlign file from a ChIP-seq BAM file for the phantompeakqualtools and MACS2 program.

For example, to generate a tagAlign file named chip_TA.tagAlign.gz from mycellchipseq.bam, execute:

~~~
sambamba view --nthreads “$NUM_THREADS" --filter 'not(unmapped or mate_is_unmapped or failed_quality_control)' mycellchipseq.bam | awk 'BEGIN{OFS="\t"} {if(and($2, 16) > 0) {print $3, ($4 - 1), ($4 - 1 + length($10)), “N”, “1000”, “-"} else {print $3, ($4 - 1), ($4 - 1 + length($10)), “N”, “1000”, “+"}}' | gzip -c > chip_TA.tagAlign.gz
~~~

You are now ready to estimate the predominant fragment lengths using cross-correlation analysis. The input file can be in tagAlign or BED format.

##### Running phantompeakqualtools

From the command line, execute the following:

1. Using Rscript, run the phantompeakqualtools program from the command line in order to use cross-correlation analysis to estimate the predominant fragment lengths. To use multiple threads with phantompeakqualtools, use the -p option. For example, to run phantompeakqualtools with $NUM_THREADS threads, execute:

~~~
Rscript phantompeakqualtools-master/run_spp.R -c=chip_TA.tagAlign.gz - p="$NUM_THREADS" -filtchr=chrM -savp=chip_TA.cc.plot.pdf -out=chip_TA.cc.qc
~~~
2. Execute the following command to write only the first value for estimated fragment length into the output file. This value (in almost all cases) is the best estimate of predominant fragment length.

~~~
sed -i -r ’s/,[^\t]+//g' chip_TA.cc.qc
~~~

The output file chip_TA.cc.qc contains NSC/RSC results in a tab-delimited file of 11 columns. The columns are filename, number of reads, estimated fragment length, strand cross-correlation at estimated fragment length, read length, strand cross correlation at read length, strand shift with minimum cross-correlation, minimum cross-correlation, normalized strand cross-correlation coefficient NSC, relative strand cross-correlation coefficient RSC, and Quality Tag.

Notably, the estimated fragment length (in column 3) can contain multiple comma-separated values. We recommend using the first value, as this value is the best estimate of predominant fragment length in almost all cases.

NSC values significantly less than 1.1 and RSC values significantly less than 1 have high background signal or low signal to noise ratios, which indicates poor quality data or low abundance of DNA-protein binding events. In our example, the ‘chip_TA.cc.plot.pdf’ output file contains the cross-correlation plot.

##### Generate fold enrichment coverage tracks using MACS2

The normalized signal track generation requires the use of MACS2 (https://github.com/taoliu/MACS/). MACS2 requires Python 2.7 (>=2.7.5) and will not work with Python 3. To install MACS2 with pip, execute the following:

##### Installing MACS2 with administrator access (all Linux distributions)

~~~
sudo pip install MACS2
~~~

##### Installing MACS2 without administrator access (all Linux distributions)

~~~
pip install --user MACS2
~~~

This section will also require bedtools, bedClip, and bedGraphToBigWig. See Box 1 for installation instructions regarding bedGraphToBigWig and bedtools. To install bedClip, execute:

~~~
wget http://hgdownload.cse.ucsc.edu/admin/exe/linux.x86_64/bedClip
chmod +x bedClip
mv bedClip ${HOME}/.local/bin/
~~~

MACS will use as input a tagAlign file and a control ChIP-seq sample tagAlign file. Use the same procedure described above to convert your control file into tagAlign format if starting with a BAM file. The MACS2 program will ultimately produce a fold-enrichment file in the bigWig format. To use MACS2 to produce this output, execute the following from the command line:

1. Create the output directory for the MACS2 results:

~~~
mkdir -p peak_output
~~~
2. Using MACS2, generate the narrow peaks and preliminary signal tracks using the tagAlign file generated directly from the BAM file. The --gsize parameter passes the effective genome size to MACS2. Using bedtools genomecov and a mappability track for a read-length of 50, we calculate the effective genome size for hg38 to be 2.8e9. MACS2 uses the effective genome size to calculate the background Poisson λ. You should always use your input ChIP-seq data’s fragment length for MACS2. For example, to generate the signal tracks for tagAlign file chip_TA.tagAlign.gz and control ChIP-seq sample tagAlign file control_TA.tagAlign.gz with a fragment length (specified with --extsize) of 220 and a p-value cutoff (specified with --pvalue) of 0.01, execute:

~~~
macs2 callpeak --treatment chip_TA.tagAlign.gz --control control_TA.tagAlign.gz --format BED --name peak_output/chip_TA --gsize 2.8e9 --pvalue 1e-2 --nomodel --extsize 220 --keep-dup all --bdg --SPMR -- shift 0
~~~ The --keep-dup all option specifies that MACS2 should keep all duplicate tags at the exact same location. The --bdg option produces peak_output/chip_TA_treat_pileup.bdg, which we use for noise removal. The --shift 0 option specifies that there should be no arbitrary shift.
3. Using MACS2, generate the final fold-enrichment signal tracks. To generate the signal tracks for our example, execute:

~~~
macs2 bdgcmp --tfile peak_output/chip_TA_treat_pileup.bdg --cfile peak_output/chip_TA_control_lambda.bdg --outdir peak_output --ofile chip_TA_FE.bdg --method FE
~~~
4. Using bedtools and bedClip, remove coordinates outside those specified in your chromosome sizes file, and generate a sorted bedGraph file. You will need a chromosome size file. This is a tab-delimited file with two columns; chromosome name (column 1), and chromosome size in base pairs (column 2). To download the chromosome sizes file, GRCh38_EBV.chrom.sizes.tsv, for hg38, see **Step 2** from the protocol. For example, to remove any coordinates in chip_TA_FE.bdg that are outside of those specified in GRCh38_EBV.chrom.sizes.tsv, and generate and sort the resulting bedGraph file, execute:

~~~
bedtools slop -i peak_output/chip_TA_FE.bdg -g GRCh38_EBV.chrom.sizes.tsv - b 0 | awk '{if ($3 != -1) print $0}' | bedClip stdin GRCh38_EBV.chrom.sizes.tsv peak_output/chip_TA.fc.signal.bedGraph
sort -k 1,1 -k2,2n chip_TA.fc.signal.bedGraph &> chip_TA.fc.signal.sorted.bedGraph
~~~
5. (Optional) Using bedGraphToBigWig, convert the resulting bedGraph file to bigWig format. Segway can directly use bedGraph files as signal tracks. However, bigWig format enables more efficient visualization on the UCSC Genome Browser of large, dense, and continuous data. For example, to convert chip_TA.fc.signal.sorted.bedGraph to bigWig format:

~~~
bedGraphToBigWig peak_output/chip_TA.fc.signal.sorted.bedGraph GRCh38_EBV.chrom.sizes.tsv peak_output/chip_TA.fc.signal.sorted.bigWig
~~~
6. (Optional) Remove intermediate files

~~~
rm -f peak_output/chip_TA_peaks.xls
rm -f peak_output/chip_TA_peaks.narrowPeak
rm -f peak_output/chip_TA_summits.bed
rm -f peak_output/chip_TA_FE.bdg
rm -f peak_output/chip_TA.fc.signal.bedGraph
rm -f peak_output/chip_TA_treat_pileup.bdg
rm -f peak_output/chip_TA_control_lambda.bdg
~~~

For problems encountered in this Box, refer to **? TROUBLESHOOTING**

#### Box 3 Downloading ChIP-seq data from ENCODE

The ENCODE Project (https://www.encodeproject.org/) provides raw and processed ChIP-seq data for transcription factors and histone modifications on its website. Locate data of interest through this website’s search or the ENCODE data matrix (https://www.encodeproject.org/matrix). The data matrix makes it easy to explore all available experiments for your cell type of interest.

For example, we want to acquire ChIP-seq data for H3K4me1 in the DOHH2 cell type. This cell type is derived from the pleural effusion of a B cell lymphoma patient. We will therefore use the search box to find it directly.

1. Open https://www.encodeproject.org/ in your browser.
2. In the search box at the upper right corner of the page, type in “DOHH2 H3K4me1 ChIP-seq” and press Enter (Figure B3.1).
3. Click on any of the results that match your preferences to visit the experiment summary page. If you plan to use data from several ChIP-seq experiments, consider that different labs may have generated the data. These labs might also use different laboratory protocols. You can read these details in the experiment summary page (Figure B2.2).
4. In the “Files” section, there are two panels for accessing raw and processed data (Figure B2.3). Each panel provides detailed information on each file. The “File type” and “Mapping assembly” columns guide you to the format you need for your analysis. If the “Biological replicate” column is empty or “1,2”, this indicates replicates are merged. Find the file type and genome assembly you need.
5. Click the download icon in the “Accession” column in the right of accession numbers to download a file immediately. Alternatively, right click on the icon and copy the link location to use with “wget" command later. In the example below, we download the signal file of H3K4me1 ChIP-seq experiment in DOHH2:

~~~
URL="https://www.encodeproject.org"
ACCESSION="ENCFF509XSM"
FORMAT="bigWig"
CELL="DOHH2"
MARK="H3K4me1"
wget “$URL/files/$ACCESSION/@@download/$ACCESSION.$FORMAT" \-O “$ACCESSION\_$CELL\_$MARK.$FORMAT"
~~~

**Figure B3.1.**
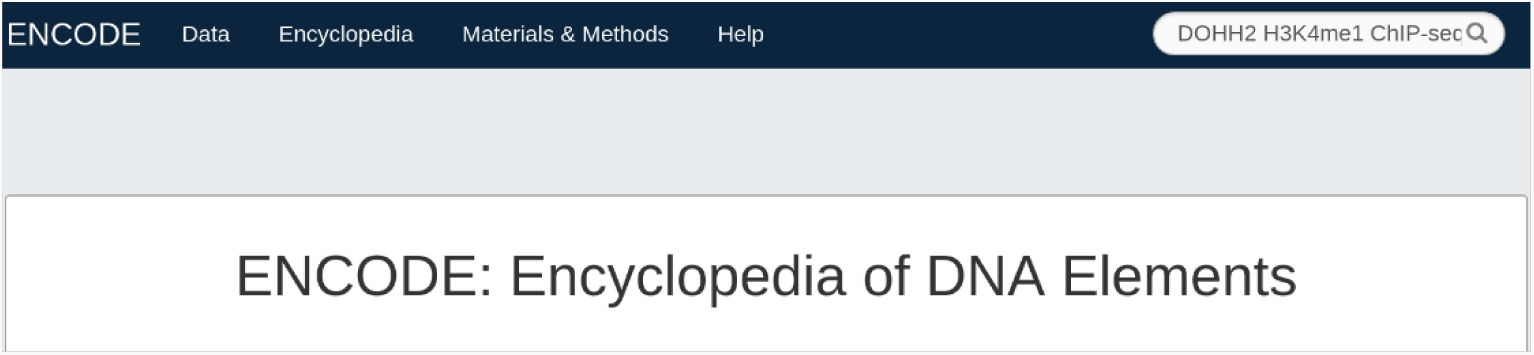
Using the ENCODE DCC search panel to find a particular experiment.

**Figure B3.2.**
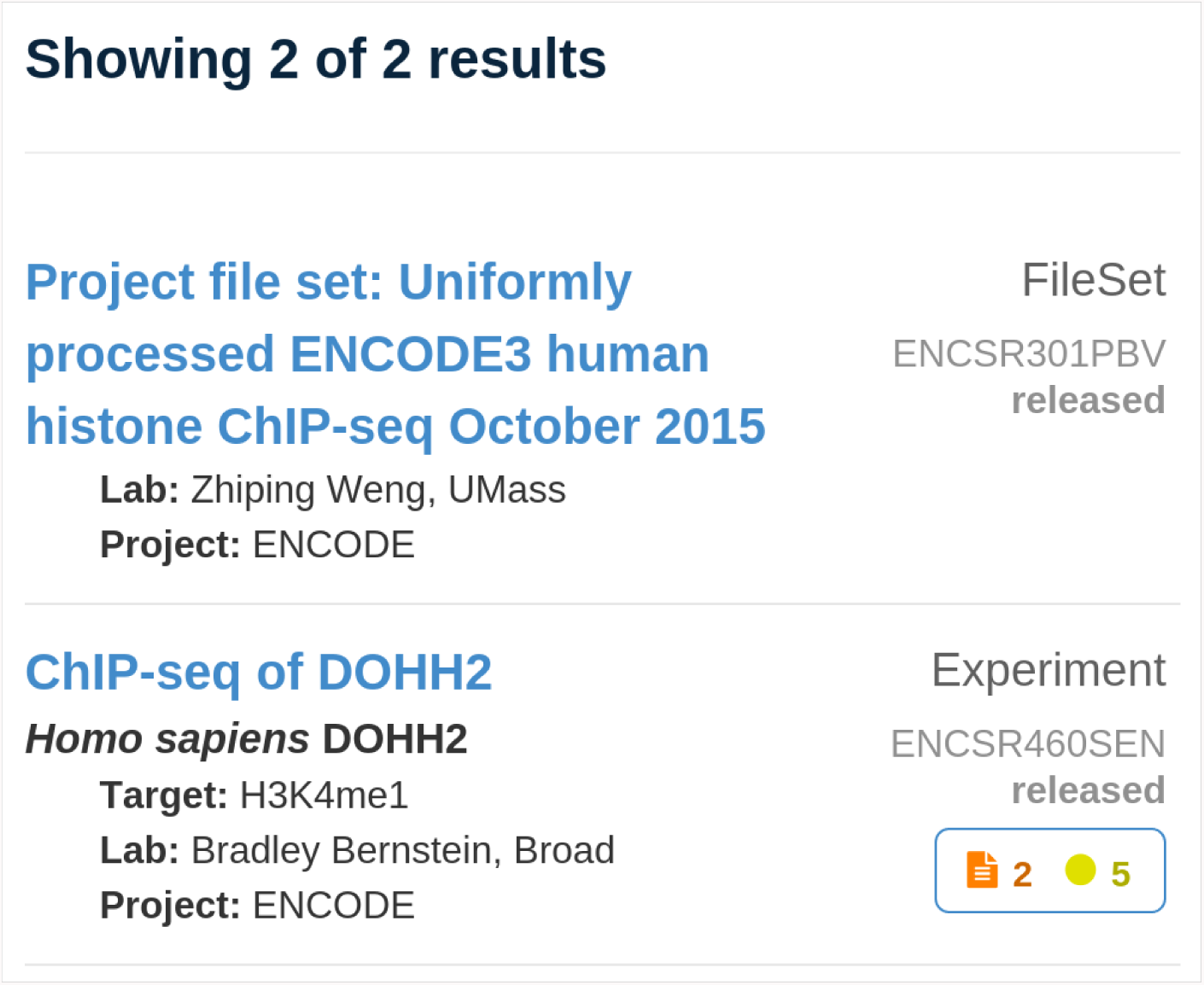
Example of the type of results you would get when using ENCODE DCC website search option.

**Figure B3.3.**
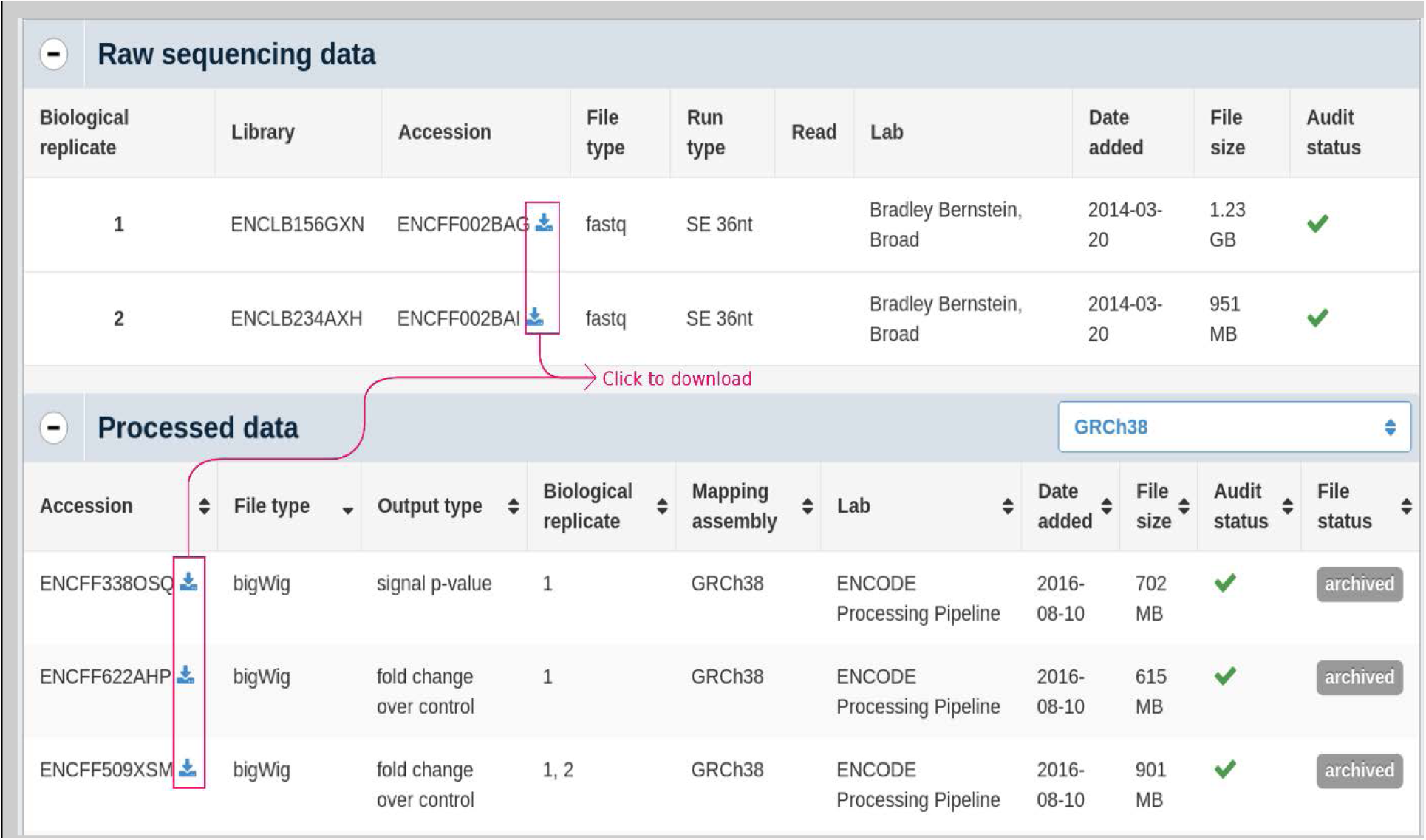
When you select the results of a search on the ENCODE DCC website, the website will direct you to a webpage on the dataset. In this webpage, you can find detailed information on aspects of the experiment. You can also download raw or processed data files (e.g. bigWig or BAM files) directly from this webpage.

ENCODE project website has a detailed application program interface (API) documentation (https://www.encodeproject.org/help/rest-api/). The ENCODE project API allows you to automate your search queries and downloads. The preferred input for Segway is the “fold change over control” bigWig signal file, because it is already processed and normalized. It is possible to generate the signal file from raw files of any ChIP-seq experiment using MACS2 (see Box 2).

For the examples in this manuscript, we download the signal files of K562 ChIP-seq datasets for H3K4me1 (ENCFF509XSM), H3K4me3 (ENCFF745GML), H3K27ac (ENCFF890NAY), H3K27me3 (ENCFF592CSV) and CTCF (ENCFF884IIL). These bigWig files average approximately 1 GiB each in size. To download all of these files, execute the following script in a bash terminal:

~~~
URL="https://www.encodeproject.org"
FORMAT=bigWig
CELL=DOHH2
MARKS=(H3K4me1 H3K4me3 H3K27ac H3K27me3 CTCF)
ACCESSIONS=(ENCFF509XSM ENCFF745GML ENCFF890NAY ENCFF592CSV ENCFF884IIL)
ITERARRAY=($(seq 0 ${#MARKS[@]}))
for i in ${ITERARRAY[@]}
do
ACCESSION=${ACCESSIONS[$i]}
MARK=${MARKS[$i]}
wget "$URL/files/$ACCESSION/@@download/$ACCESSION.$FORMAT" \
-O "$ACCESSION.$CELL.$MARK.$FORMAT"
done
~~~

#### Box 4 Quality control for ChIP-seq data

For Segway to produce high-quality segmentations, input chromatin immunoprecipitation sequencing (ChIP-seq) data must have sufficient quality. Quality control methods for ChIP-seq data can ensure this.

Several technical factors affect the quality of ChIP-seq data.^42^ Often, most important is the specificity of the antibody employed. Also key is using a sufficient number of cells and appropriate controls. Fragmentation, library construction, and sequencing protocols used also influence data quality.^42^ Finally, the alignment software used influences signal mapping quality.

This box explains how to compute different quality control metrics using FastQC,^43^ ChIPQC,^44,45^ and NGS-QC.^46–48^ We provide general guidelines, which provide a clear means of assessing data quality. There are, however, a number of different ways of assessing quality and many methods depend greatly upon the particulars of the assessed experiment. There is not yet a consensus on a general and optimal means of assessing ChIP-seq data quality and we suggest that interested readers consult some of the broad literature on this topic.^38,39,44,46,49–52^

##### Installation of QC tools and dependencies

FastQC^43^: follow instructions at http://www.bioinformatics.babraham.ac.uk/projects/fastqc/INSTALL.txt.

Sambamba^41^: follow instructions at http://lomereiter.github.io/sambamba or our instructions in Box 2.

ChIPQC:^44,45^

1. Install: R^53^ 3.3.0
2. Install Bioconductor^54^ 3.3 and ChIPQC version 1.8.8, by executing the following within the R environment:

~~~
source(“http://bioconductor.org/biocLite.R”)
biocLite() # install Bioconductor
biocLite(“ChIPQC”) # install ChIPQC
~~~

##### Sequence data quality control with FastQC

FastQC^43^ reports on potentially problematic aspects of the sequencing itself. This includes base qualities, G+C bias, systematic overrepresentation of sequences, and several other metrics. To run FastQC on BAM files, such as the ENCODE CTCF ChIP-seq samples, whose signal files were downloaded in Box 3, execute:

~~~
fastqc ENCFF863PSQ.bam ENCFF092CZO.bam
~~~

One useful metric is library complexity—a measure of the number of distinct molecules in the library. Low complexity often results in repeated sequencing of duplicates, yielding little information.^55^ You can estimate library complexity from FastQC’s duplicate sequence plot. More detailed analyses can be performed, if necessary, using preseq.^56^ A large fraction of duplicate sequences is often a result of insufficient sequence diversity, which can suggest an inherent experimental bottleneck, such as an insufficient number of input cells.^50,55^

Additional considerations include checking the quality of read mapping, the proportion of reads aligning to a genomic position, the number passing a mapping quality threshold, and the abundance of duplicate reads.

##### Assessing ChIP-seq data quality

To assess ChIP-seq data quality; you should perform overall assessment of both ChIP-seq reads and the effects of random sub-sampling, when possible. You should use frequency of reads in peaks (FRiP) with ChIP-seq peak calls, but we do not focus on this approach since Segway operates directly upon signal. It is useful to assess the consistency across ChIP-seq replicates, such as via an irreproducible discovery rate^57^. ENCODE conducts this analysis for all of its ChIP-seq datasets.^39^ This is currently non-trivial, however, to use in one’s own workflow.

##### Testing ChIP-seq quality with ChIPQC

ChIPQC^44,45^ is an R^53^ Bioconductor^54^ package that evaluates metrics of mapping, filtering, and duplication rates, as well as ChIP-seq signal distribution and structure. While ChIPQC usually also computes FRiP, from called peaks, here we will run it using only signal data.

1. Define or use the previously exported NUM_THREADS environment variable.
2. Determine the maximum memory for ChIPQC to use in GiB:

~~~
MAX_M_USE=$(($(free -g | head -3 | tail -1 | tr -s ' ' | cut -d ' ' -f 4) -1))
~~~

-->--> On a cluster, use instead 1 GiB less than the amount allocated to your job.
3. Create a sorted, indexed, and duplicate marked BAM file, using Sambamba^41^. For example to do so for the ENCFF863PSQ.bam, run the following:

~~~
a. sambamba sort --memory-limit “${MAX_M_USE}GiB" --nthreads "$NUM_THREADS" --compression-level 0 --out “${TMPDIR:-/tmp}/ENCFF863PSQ.sorted.bam" ENCFF863PSQ.bam
b. sambamba markdup --nthreads “$NUM_THREADS" --compression-level 9 --tmpdir="$TMPDIR" “${TMPDIR:-/tmp}/ENCFF863PSQ.sorted.bam"/dev/stdout | tee ENCFF863PSQ.sorted.markeddup.bam | sambamba index --nthreads “$NUM_THREADS" /dev/stdin
ENCFF863PSQ.sorted.markeddup.bam.bai
rm -f “${TMPDIR:-/tmp}"/ENCFF863PSQ.sorted.bam*
~~~
4. Create an experiment description file for ChIPQC. This file describes the ChIP-seq samples that ChIPQC will analyze. ChIPQC operates on each input file, which might be a single technical replicate, with pooled sequencing lanes, or a single biological replicate, merged from multiple technical replicates. The content of a single unit of quality assessment—a single file—depends upon your experimental setup and downstream experimental goals. Each row corresponds to one ChIP-seq file, while each column describes the data associated with that file. We highlight a common use-case; refer to the package documentation for a detailed description of all available fields. You should specify the following columns: SampleID, Tissue, Factor, Replicate, bamReads, ControlID, bamControl, and Peaks. These fields act merely as annotations of your data and do not alter ChIPQC’s operation, with two exceptions. Two fields must contain valid file paths: bamReads, which must contain the full path to the above sorted and duplicate marked BAM file of the ChIP-seq experiment. The other field, bamControl, must contain the full path to a corresponding set of control reads, in a sorted and duplicate marked BAM, such as from an input (antibody-free) experiment. In this case, without peak calls, set Peaks to NA. This will cause ChIPQC to compute all metrics that do not depend upon a peak set. If you have a peak file, set Peaks to the file name and additionally specify a PeakFormat column, if the file is not in BED format. Even if you are only analyzing a single replicate, you must still specify the Replicate column. You may set it to 1 in this case. Name this file QCexperiment.csv and delimit its columns with tabs. For example, a single replicate may result in a file like this:

~~~
SampleID Tissue Factor Replicate bamReads ControlID
bamControl Peaks
ENCFF863PSQ DOHH2 CTCF 1 ENCFF863PSQ.sorted.markeddup.bam
ENCFF631ENA ENCFF631ENA.sorted.markeddup.bam NA
~~~
5. Execute the following within the R environment:

~~~
library(ChIPQC)
samples <- read.delim(“QCexperiment.csv”, stringsAsFactors=FALSE)
experiment <- ChIPQC(samples, annotation="hg38”)
~~~ Specify the assembly employed via the annotation parameter. We used GRCh38/hg38 above, but you can also, for example, use GRCh37/hg19 instead via annotation="hg19".
6. Generate the output report, summary, and plots of interest by executing in the R environment:

~~~
# disable faceting when using only a single sample
facet <- ifelse(nrow(samples) > 1, TRUE, FALSE)
write.table(QCmetrics(experiment), file='QCmetrics.csv')
ChIPQCreport(experiment, facet=facet)
pdf('plots.pdf')
plotCoverageHist(experiment, facet=facet)
plotCC(experiment)
plotSSD(experiment, facet=facet)
if (!all(is.na(samples$Peaks))) {
  plotPeakProfile(experiment)
}
dev.off()
~~~

The ChIPQC documentation contains additional details, including other available plots.^45^

##### ? TROUBLESHOOTING

###### Assessing read mapping quality

Verify that ChIP-seq results have a substantial portion of uniquely mapped reads, without an unexpectedly high proportion of reads filtered out due to insufficient quality. Generally, at least 50% of reads in an experiment should map uniquely, with lower values expected for input data, which lacks a targeting antibody. This varies greatly, however, and depends on the species analyzed.^50,51^ In human and mouse ChIP-seq samples, expect over 70% of reads to map uniquely.^50,51^ You should also assess the duplication rate—a ratio of unique to total reads. This varies greatly, but you should expect for it to be much less in control samples and it should generally not exceed 50%.^45^

###### Assessing ChIP-seq signal distribution

You should also assess the read cross-correlation, as a metric of ChIP-seq data quality. The ChIPQC cross-correlation plot should have a clear peak at the fragment length in successfully-enriched samples.^45^ The normalized strand cross-correlation coefficient within the QC metrics file should be greater than 1.05, while the relative strand cross-correlation coefficient should be greater than 0.8.^39^ Bailey et al.^50^ describes how to use and interpret this metric further (Box 2 of Bailey et al.^50^).

Additionally, evaluate ChIPQC’s coverage histograms and their standardized standard deviation (SSD), normalized to read depth. Expect the coverage histogram to have a non-negligible “tail” and for SSD values to be greater than 1, generally above 1.5. Expect controls to have SSD values of around 1. Control SSDs greater than 1 might indicate aberrant enrichment.^45^

##### Subsampling to assess ChIP-seq quality with NGS-QC

Random sub-sampling on ChIP-seq profiles provides a means of computing quality metrics that do not depend upon peak calling. Importantly, such metrics are comparable between sharp (transcription factors) and broad (histone modifications) peaks,^46^ both of which are frequently used together by Segway. The Next Generation Sequencing Quality Control Generator (NGS-QC)^46–48^ provides these metrics for a wide array of ChIP-seq datasets. It uses measurements of global deviations of random read subsets with respect to the full set of aligned reads to assign quality scores.^48^ NGS-QC randomly subsamples 90%, 70%, and 50% of reads and counts reads in subsampled and full datasets in 500–base-pair bins. NGS-QC measures the variance from the expectation of the same percentage decrease in read counts. It quantifies the proportion of these bins that are below 2.5%, 5%, or 10% of this expected fraction, and each threshold forms a component of the quality score. These scores form labels for ChIP-seq data quality from A–D, for each threshold. Therefore, the highest rating is “AAA”, while the lowest is “DDD”.^48^ NGS-QC has pre-computed quality metrics for many public datasets, including some ENCODE data. Consult these if available. Otherwise, compare your data to them, as outlined below.

Use NGS-QC via its Galaxy^58^ instance, as follows:

1. Navigate to: http://galaxy.ngs-qc.org/
2. Create an account.
3. Upload BAM files, using FTP.

a. From the directory containing the BAM files to upload on the command line, run ftp galaxy.ngs-qc.org
b. Login using the previously created credentials.
c. Execute:

i. prompt (disables confirmation of each individual upload)
ii. mput <files>
iii. exit
d. Navigate to the Galaxy instance, and select “Get Data” > “Upload file” from the left panel.
e. Select the files uploaded via FTP, using the “Choose FTP file” option. Access the NGS-QC command under “NGS-QC” > “NGS-QC Generator“. Fill in the provided form to assess the uploaded data, following the NGS-QC guidelines (http://www.ngs-qc.org/tutorial.php#part2). We recommend selecting three replicates of the QC computation, to mitigate against sampling bias. Specify the genome to which the uploaded data was aligned, from the available list. At this time, GRCh38/hg38 is not available. For the other parameters, use the defaults.
4. Compare against existing public datasets, selecting the target of the ChIP-seq experiment at the bottom of the “NGS-QC Generator” interface, when it lists your target.
5. Once Galaxy indicates that the tasks have completed (turning green), select the “View Data” button for the task starting with “Results”. Then inspect each replicate, especially its quality rating and percent uniquely mapped reads. Evaluate them using the <u>criteria above</u>.

Mendoza-Parra et al.^47^ and the NGS-QC tutorial (http://www.ngs-qc.org/tutorial.php#part4) provide details on how to interpret the results.

#### Box 5 Accelerating Segway using a compute cluster

We designed Segway to run on a large variety of cluster systems. Segway uses the Distributed Resource Management and Application API^59^ (DRMAA) version 1 for submitting its jobs to a cluster. This interface has been implemented for a variety of cluster systems including Grid Engine^60^ (GE), Condor^61^, Portable Batch System^62^ (PBS / Torque), and Platform Load Sharing Facility^63^ (LSF).

##### Installing DRMAA

We recommend that a cluster administrator installs DRMAA. With the exception of Grid Engine, which has DRMAA installed by default, each cluster system has its own specific installation procedure. The general steps for installing DRMAA are as follows:

1. Download and build the DRMAA implementation for your cluster system: **Grid Engine** Not applicable: installed by default **Torque or PBS Pro** https://sourceforge.net/projects/pbspro-drmaa/ **Platform Load Sharing Facility** https://sourceforge.net/projects/lsf-drmaa/
2. Set the environment variable DRMAA_LIBRARY_PATH to the full name and path of the DRMAA library. For example, if you have GE installed in /sge/lib/linux-x64 execute:

~~~
export DRMAA_LIBRARY_PATH="/sge/lib/linux-x64/libdrmaa.so"
~~~

##### Compute cluster settings in Segway

Some cluster configurations require submitted jobs to have specific settings. Segway can specify cluster specific settings with the --cluster-opt option. This option passes on what would normally be set as options from your native job submission command (such as qsub from Grid Engine or Torque). For example, for Segway to submit jobs to a Grid Engine queue named bioinformatics use the option:

~~~
--cluster-opt="-q bioinformatics"
~~~

##### Verifying job submission by Segway

When Segway submits jobs, it logs each submission to the console. If the log entry starts with “queued” then DRMAA is running successfully. If the log entry starts with “running locally”, then Segway is not submitting the jobs to the cluster and they are instead running “locally” on the same machine that contains the Segway process.

#### Box 6 Comparing with Published Segway Annotations

The Segway website (<u>segway.hoffmanlab.org</u>) provides access to publicly available genome segmentations. Comparing your segmentation data with previous annotations will help you to assign biological meaning to the numbered labels for each genomic region (for example TSS, promoter, or enhancer). One strategy for making these comparisons is to visualize your segmentation alongside a published segmentation in the UCSC Genome Browser, and to compare your labeled regions to annotated labels. Another strategy is to determine the similarity of intervals associated with a numbered label in your segmentation to regions with a mnemonic label from a published segmentation.

In this example, we will compare the DOHH2 segmentation to a published segmentation of the GM12878 cell line from the Ensembl Regulatory Build, which partitions the genome into regulatory regions (for example, active promoter, polycomb repressed, or distal enhancer) for comparison in the UCSC Genome Browser. DOHH2 is a B-cell myeloma line and GM12878 is lymphoblastic B-cell line. We will use the track hub outlined in this protocol for visualization. The Regulatory Build segmentation employs the key shown below for identification of the regulatory elements represented by the colored and named labels^64^.

**Figure B6.1:**
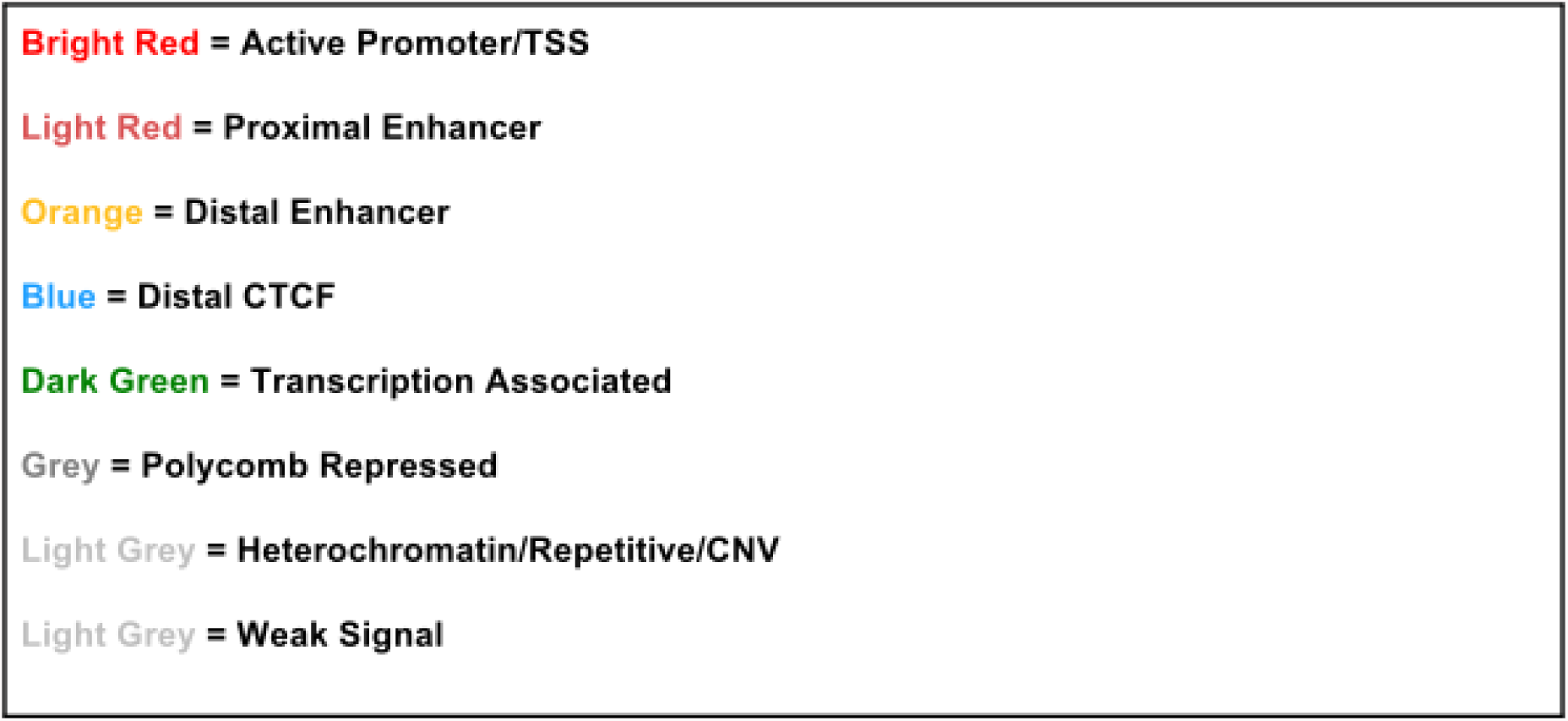
Classification of regulatory elements for labels in the Ensembl Regulatory Build.

##### Visualization and comparison of published segmentation

You can visualize the published segmentation through the UCSC Genome Browser Track Hub portal or by downloading the segmentation file and adding the file to your personal track hub.

Load Ensembl Regulatory Segmentation from the UCSC Genome Browser:

1. Navigate to the UCSC Genome Browser (https://genome.ucsc.edu/). Select “My Data” > “Track Hubs.”
2. Under “Public Hubs”, connect the “Ensembl Regulatory Build.”
3. After loading the hub, open the genome browser to the hg38 assembly and the regulatory track hub will appear in your track hub.
4. To visualize only the GM12878 segmentation, right click on the segmentation in the browser window and click “Configure Cell Type Activity”
5. Select only the GM12878 track and click “Submit.”

**Figure B6.2:**
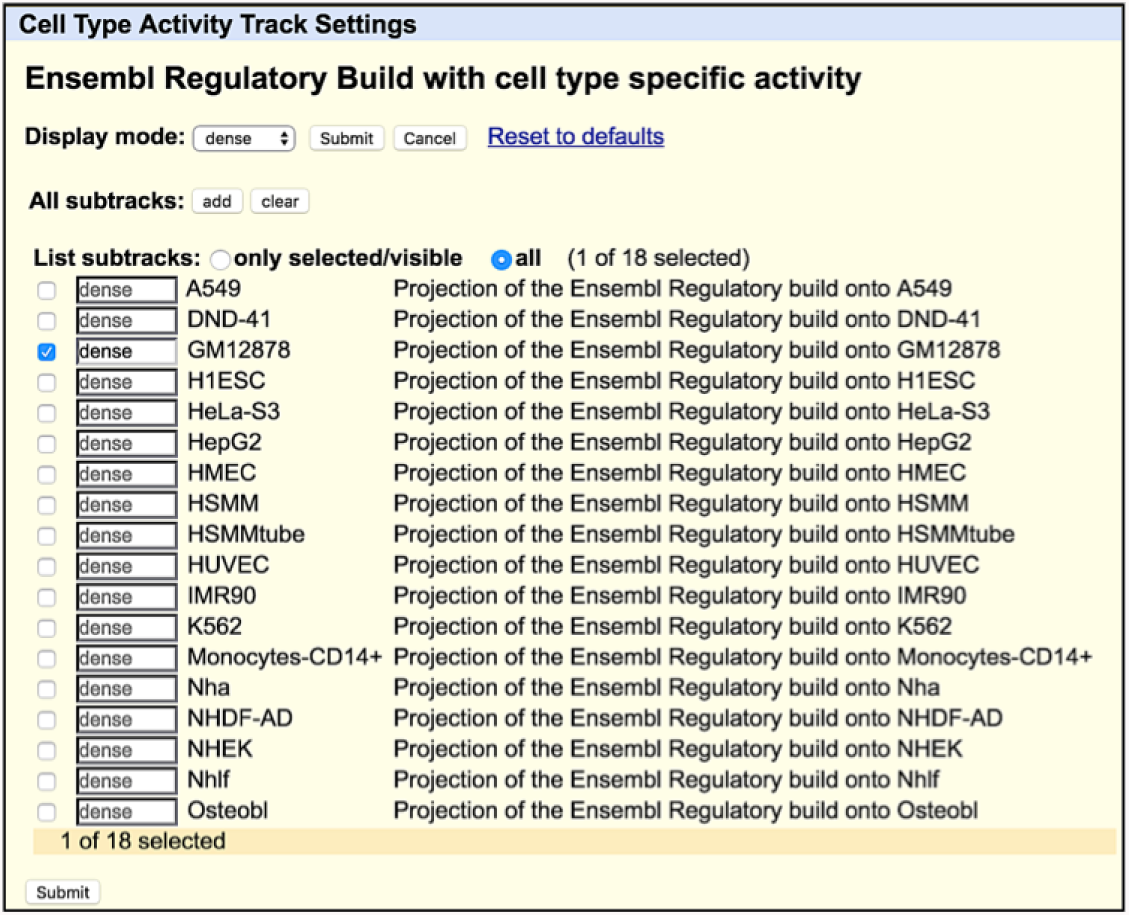
Loading cell type specific segmentation into the UCSC Genome Browser session

Add the Ensembl Regulatory Segmentation to your track hub:

1. Open segway.hoffmanlab.org in your browser.
2. Click on the link “Ensembl Regulatory Build for GRCh38 (hg38).”
3. Click the link called “segmentations/” and in the files section of the new page copy the link location for the GM12878 and use the “wget” command to download the bigBed file for this cell line.
4. Add the regulatory segmentation file to your “trackDb.txt” file using the same format described for the DOHH2 segmentation file.
5. Use the “rsync” command to upload the GM12878.bb file to the publically available server used for your Track Hub.
6. Open your track hub in the UCSC Genome Browser. The regulatory segmentation and your segmentation file should appear in the browser window.

In the example below, we examine the *MEN1* gene locus within the DOHH2 segmentation and the GM12878 published regulatory segmentation. You may also include the signal tracks loaded into the track hub when visualizing the segmentations, which may provide additional information about genomic features captured by segments and labels. *MEN1* codes for menin protein, which acts as a tumor suppressor and is dysregulated in many cancers^23^ (Figure B6.3).

**Figure B6.3:**
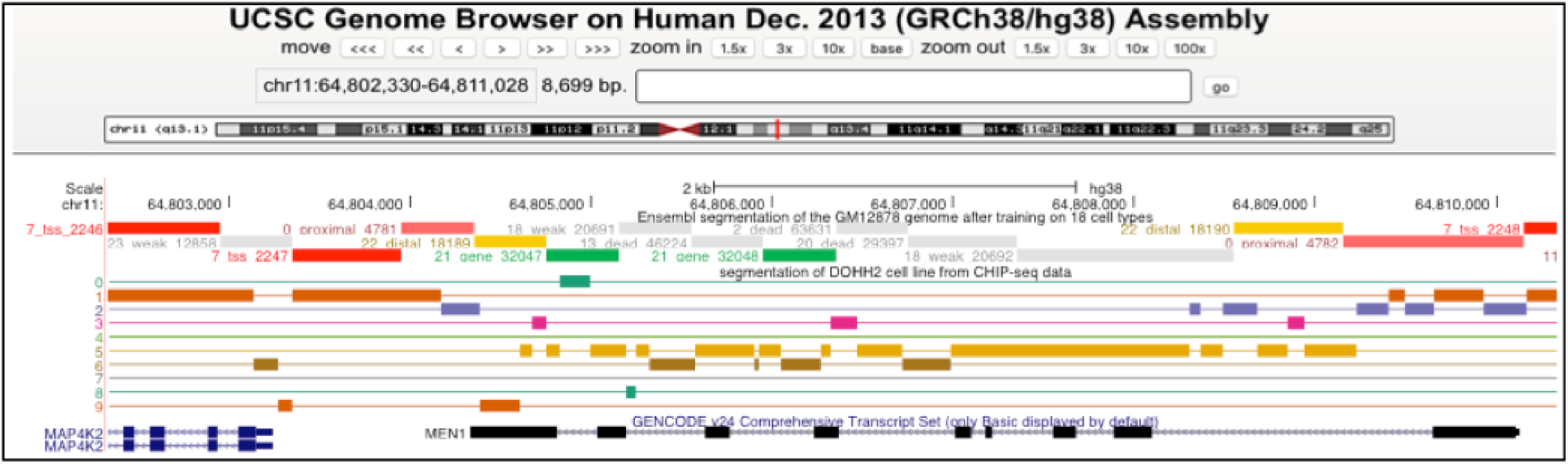
Comparison of Ensemble Regulatory segmentation for GM12878 cell line and DOHH2 segmentation at the *MEN1* locus.

The regulatory build segments identified as TSS (red) and proximal enhancer (light red) regions align with the first exon of MEN1 and a neighboring gene MAP4K2. The DOHH2 segmentation produced segments in label 1 that overlap the TSS and proximal enhancer segments of the regulatory build and the first exons of the visualized genes. Visual comparison at other genes may identify label 1 segments as correlated to proximal enhancers, promoters and TSS.

Use the “zoom in” tool on the browser for a closer view of the segments.

**Figure B6.4:**
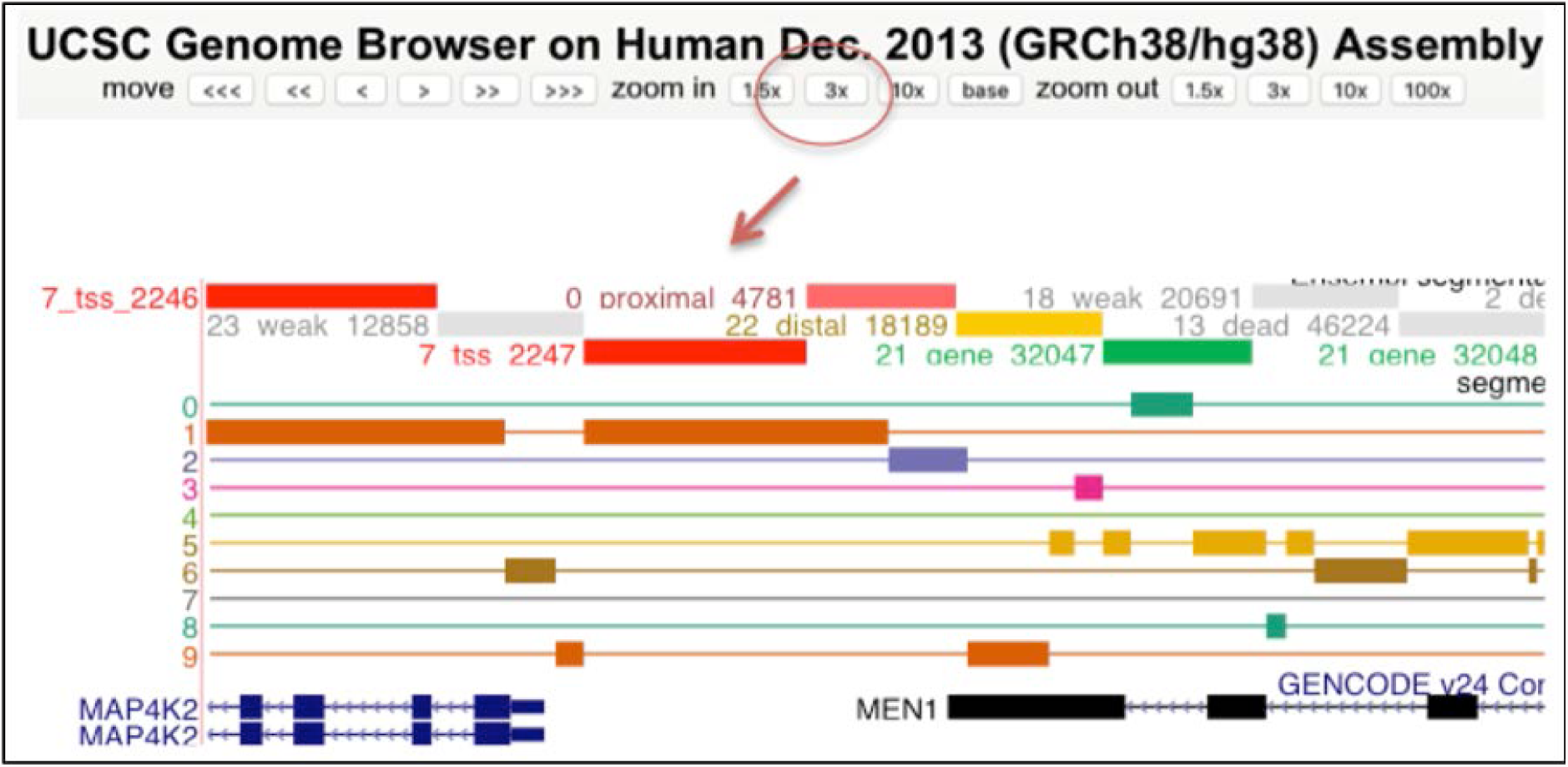
Identification of enhancer regions at the *MEN1* locus. The regulatory segmentation also identifies 2 distal enhancer regions at the *MEN1* gene. This figure shows segments in labels 3, 5, and 9 that overlap with 1 of these identified enhancer regions. This type of visual comparison at other genes may identify one of these labels as highly correlated to overlap CTCF binding sites or enhancer regions identified by the regulatory segmentation.

Reorder the segmentation and signal tracks in the browser to facilitate visual comparison by clicking on the left-hand side of the track and dragging up or down to a new position. Additionally, you can adjust the configuration settings for each track by right clicking and selecting “Configure.” This allows you to manipulate settings, such as the density of the display, axis display height, and color display (**Figure B6.5**). In the configuration settings, you can also use subsets of the regulatory segmentation to display in the browser. For example, you can display a summary track from the regulatory segmentation that only includes segments identifying transcription factor binding sites from all cell types used in the Regulatory Build.

**Figure B6.5:**
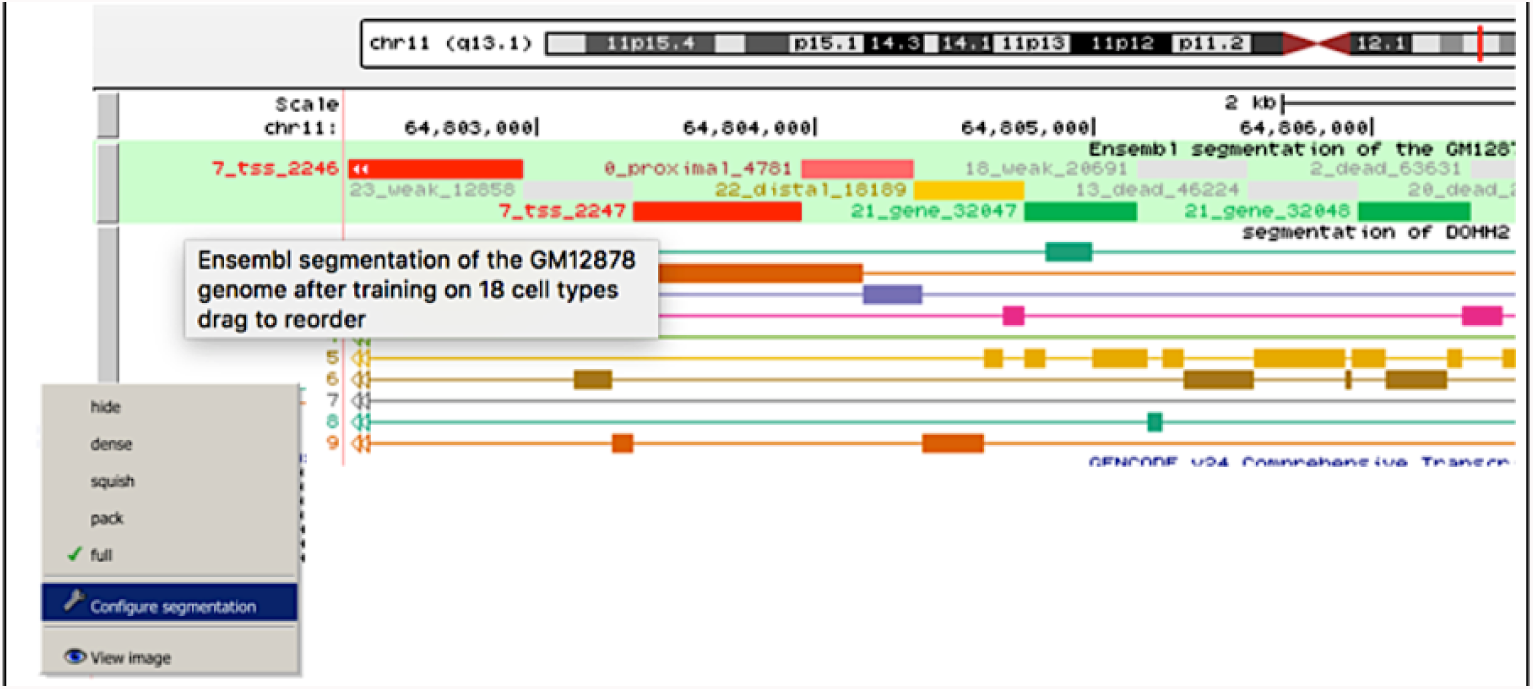
Configuring tracks in track hub for comparison of segmentations.

##### Compare segmentation files on the command line

You can also compare your segmentation to a published segmentation by identifying overlapping genomic regions between the two files and calculating the percent of overlapping regulatory element segments within each of the labels in your segmentation file. This can be accomplished using the bedtools intersect command. This will interrogate how often regions within a label from your segmentation overlap with annotated regulatory labels (e.g. tss, ctcf, dead) from the published segmentation. This will provide information about which regulatory elements may be enriched in a label from your segmentation, which will help you to assign a biologically meaningful label.

In this example, we compare the published GM12878 segmentation to the DOHH2 segmentation produced from the protocol. Bedtools enables common genomic analyses on the command line. Box 1 contains instructions for installing bedtools. Additionally, install the program “bigBedToBed” (http://hgdownload.cse.ucsc.edu/admin/exe/linux.x86_64/bigBedToBed) by executing the same commands used to install bedToBigBed in Box 1. If you did not previously download the GM12878 segmentation, follow the directions outlined above to obtain this file now.

1. Convert the published segmentation file in bigBed file format to bed file format. Execute:

~~~
bigBedToBed GM12878.bb GM12878.regulatory.bed
~~~
2. Use bedtools to intersect the 2 segmentation BED files:

~~~
bedtools intersect -a segway.bed -b GM12878.regulatory.bed -wa -wb > intersect.bed
~~~
3. Filter the resulting bed file from the intersection command to identify, count, and sort the number of times a regulatory element from the published segmentation overlaps a segment in a specific label. For this example, we check the published segmentation overlapping label 9.

~~~
awk 'BEGIN{FS=OFS="\t"} ($4 == “9”) {print $4, $13}' intersect.bed | sed
’s/_/\t/g' | cut -f3 | sort | uniq -c | sort -k1n > label.9.txt
~~~
4. Calculate the percentage of times a regulatory element is counted within a specific label

~~~
awk 'NR==FNR{a = a + $1; next} {c = ($1/a)*100; print $1, $2, c}' label.9.txt label.9.txt > comp.label9.txt
~~~

These commands will generate a plain text (txt) file as shown below (**Figure B6.6**). Label 9 overlaps 40% of segments in the regulator segmentation with the label “dead” and overlaps with 20% of overlapping regions with the label “ctcf,” which is a transcription factor binding site. As most of the genome is untranscribed, a high percentage of “dead” regions within a label is not unexpected. The relatively high percentage of regions identified as “ctcf” elements suggests that label 9 may contain many segments associated with transcription factor binding sites, such as enhancers and promoters. In a visual comparison of the DOHH2 segmentation at the Menin locus to the GM12878 segmentation, we observed a segment in label 9 that overlapped with a distal enhancer element. In label 1, proximal enhancer elements, distal enhancer elements, non-transcribed intergenic regions, and TSS compose the majority of elements from the regulatory segmentation that overlap with regions in the DOHH2 labels. We also observed with pattern with visual comparison of our segmentation files in the browser. Performing this analysis on all your labels may provide additional insights into patterns observed upon visual comparison.

**Figure B6.6:**
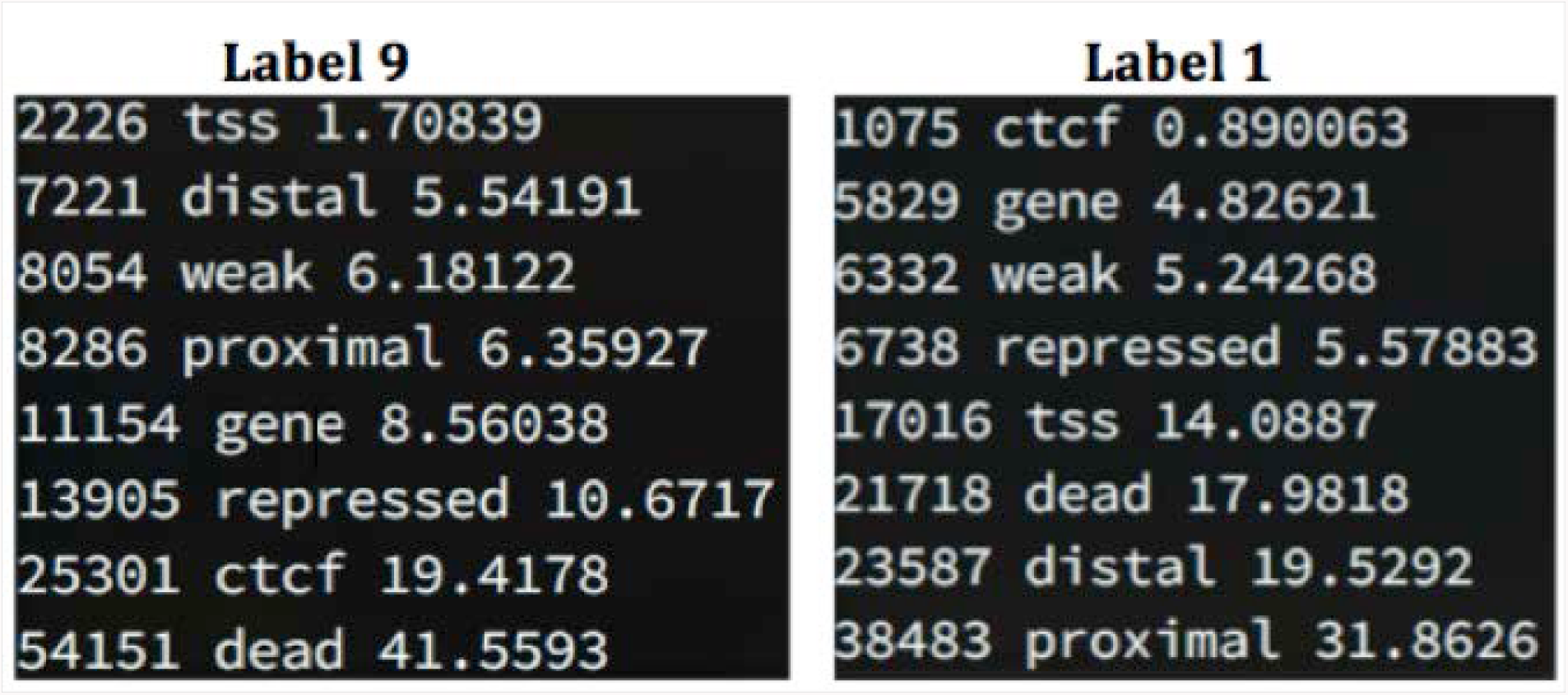
Percentage of regulatory elements from Ensembl segmentation that overlap with labels 1 and 9 in the DOHH2 segmentation.

